# Outer membrane proteins in *Mycobacterium tuberculosis* and its potential role in small molecule permeation

**DOI:** 10.1101/2023.09.16.557957

**Authors:** Aseem Palande, Saniya Patil, Anjali Veeram, Soumya Swastik Sahoo, M Balaji, Jeetender Chugh, Raju Mukherjee

## Abstract

Increased resistance to current anti-mycobacterial and a potential bias towards relatively hydrophobic chemical entities highlight an urgent need to understand how current anti-TB drugs enter the tubercle bacilli. While inner membrane proteins are well-studied, how small molecules cross the impenetrable outer membrane remains unknown. Here we employed mass spectrometry-based proteomics to show that octyl-*β*-glucopyranoside selectively extracts the outer membrane proteins of *Mycobacterium tuberculosis*. Differentially expressed proteins between nutrient replete and depleted conditions were enriched to identify proteins involved in nutrient uptake. We demonstrate cell surface localization of seven new proteins using immunofluorescence and show that overexpression of the proteins LpqY and ProX leads to hypersensitivity towards streptomycin, while expression of SubI, FecB2, and Rv0999 exhibited higher membrane permeability, assessed through EtBr accumulation assay. Further, proton NMR metabolomics suggests the role of four outer membrane proteins in glycerol uptake. This study identifies several outer membrane proteins that are involved in the permeation of small hydrophilic molecules and are potential targets for enhancing uptake and efficacy of anti-TB drugs.

## Introduction

*Mycobacterium tuberculosis* is one of the oldest human pathogens, having survived for nearly 70,000 years, and was responsible for the highest number of fatalities caused by a single infectious agent prior to the emergence of COVID-19 (1). Since the discovery of bedaquiline in 2005, the drug-discovery pipeline for Tuberculosis (2) has emerged very promising. Two decades of collaborative screening campaign have resulted in several potential new chemical entities at different stages of development. Few common features that have emerged from the successful phenotypic screens indicate a potential bias towards identifying a high number of membrane-associated protein as new targets for compounds which are relatively hydrophobic (i.e. with a higher n-octanol/water partition coefficient, log*P*) (Table 1)(3). In spite of representing diverse chemical class multiple pharmacophores appear to target the same set of protein. This has led to the classification of DprE1, MmpL3, Pks13 and QcrB as “promiscuous” drug targets (4).

**Table 1:** log*P* values of some approved and candidate anti-TB drugs (3)

| First Line TB Drugs |  | Second line TB drugs |  | Candidate TB drugs under development |  |
| --- | --- | --- | --- | --- | --- |
| Pyrazinamide (PZA) | -1.23 | Linezolid | 0.64 | Bedaquiline | 7.13 |
| Ethambutol (ETH) | -0.06 | Moxifloxacin | -0.5 | PA-824 | 4.14 |
| Isoniazid (INH) | -0.67 | Levofloxacin | 0.029 | Q203 | 5.6 |
| Rifampicin (RIF) | 2.77 | Capreomycin | -9.75 | Delamanid | 6.14 |
|  |  | Kanamycin | -7.06 | SQ109 | 4.64 |
|  |  | Streptomycin | -7.65 | BM212 | 4.75 |
|  |  | D-cycloserine | -2.42 | AU1235 | 5.1 |
|  |  | Ethionamide | 1.33 | TCA1 | 3.2 |

Mycobacterial outer membranes are highly lipid-rich, with both covalently bound and free lipids, which explains why phenotypic screening tends to favour the identification of hydrophobic compounds. The cell envelope is a unique amalgamation of a thick peptidoglycan characteristic of Gram-positive character and an outer membrane, characteristic of Gram-negative bacteria (Fig 1). The peptidoglycan is covalently attached to the arabinogalactan, which is further decorated by chains of mycolic acids forming the outer membrane. It is this lipid bilayer that confers intrinsic resistance to *M. tuberculosis* against drugs by providing a permeability barrier for both hydrophilic and hydrophobic compounds and harbours virulence factors that make it a successful pathogen. Studies on the dynamic nature of the mycobacterial outer membrane, following the incorporation of fluorescent trehalose molecules during cell wall biogenesis revealed a lack of fluidity in the outer leaflet and an extremely low rate of lateral diffusion of trehalose glycolipids (5). Outer membrane proteins (OMP) interspersed in this mycolic acid layer, allows small molecule nutrients and possibly drugs to permeate, though the process of uptake is not clearly known (6). Despite their critical role in nutrient uptake, host-pathogen interactions, and secretion of virulence factors, not many mycobacterial OMPs were studied, due to the difficulty in separating the mycobacterial inner and outer membrane. Attempts at identifying OMPs through homology-based search have been insufficient in case of mycobacteria, compared to the widely studied OMPs in Gram-negatives.

**Figure 1:**
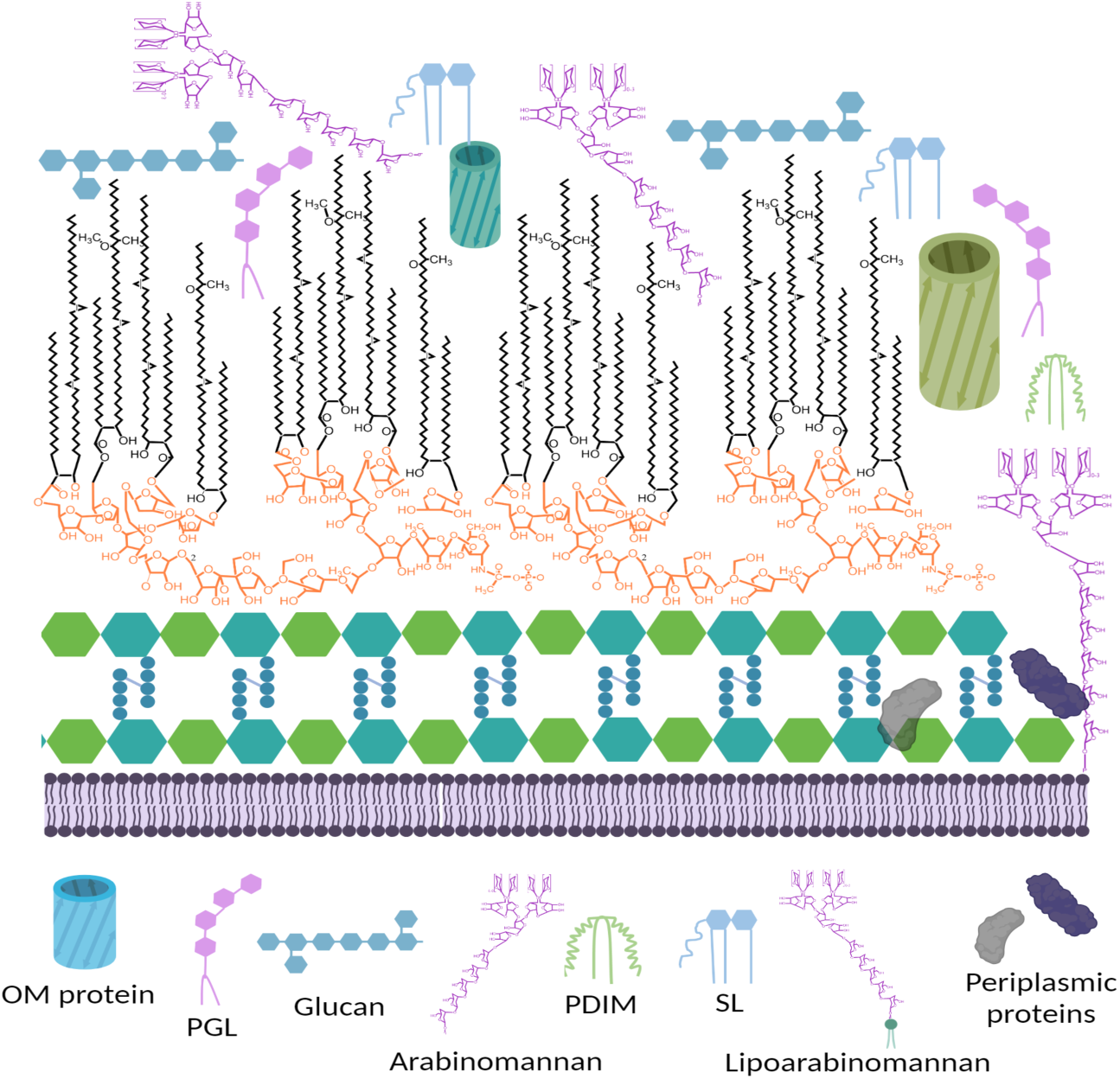
Mycobacterial cell wall. The schematic shows the complex composition of the mycobacterial surface with a focus on the outer membrane. The inner membrane with its phospholipid bilayer is surrounded by a peptidoglycan with an unconventional 3→3 crosslink between the tetrapeptides during stress conditions and stationary phase, in addition to the usual 3→4 linkage. The peptidoglycan is covalently linked to the arabinogalactan, which is decorated by long chains of covalently linked mycolic acids. This second lipid bilayer of 60-90 carbon atoms is called the outer membrane which is interspersed with phenolic glycolipids (PGL), sulfolipids (SL), phthiocerol dimycocerosates (PDIM), glucans and arabinomannans.

Water-filled channels called porins are involved in the uptake of small and large hydrophilic nutrients (7). They are goblet shaped oligomeric proteins with anti-parallel *β*-sheets first reported in Gram-negative organism and were implicated for permeation of penicillin, carbapenems, cephalosporins and fluoroquinolones (8). The polar residues inside the lumen are selective with respect to the shape or charge present on the hydrophilic substrates (9). Energy-independent passive diffusion across the outer membrane has been suggested as the mode of entry for many antibiotics in mycobacteria. Though much lower than Gram-negatives, pore forming proteins have also been observed in the fast-growing *M. smegmatis* whose genome codes for 4 porin paralogues. *mspA* and *mspC* are constitutively expressed while *mspB* and *mspD* were detected only upon depletion of the major porin *mspA*. Studies in *M. smegmatis* proposed porin-mediated uptake of several classes of approved drugs. Reduced transport of hydrophobic fluoroquinolones were observed upon treating *M. tuberculosis* with polyamines, substrates that are known to close porin channels in *E. coli* (10). Proposed modes of uptake for individual antibiotics are summarized in Table 2.

**Table 2:** Mode of entry of antibiotics through the mycobacterial outer membrane.

| Antibiotics | Property | Proposed mode of uptake | Reference |
| --- | --- | --- | --- |
| Ampicillin<br>Cephaloridine | Zwitterionic | Porin mediated | (11,12) |
| Chloramphenicol<br>Isoniazid<br>Ethambutol<br>Tetracycline | Small hydrophilic | Unspecific passive diffusion | (11,13,14) |
| Kanamycin<br>Vancomycin | Large hydrophilic | Probably Porin mediated or by self-promoted as in Gram-negatives | (15) |
| Erythromycin<br>Rifampicin | Large hydrophobic | Passive | (16) |
| Ciprofloxacin<br>Moxifloxacin<br>Levofloxacin<br>Norfloxacin | Fluoroquinolones | Porin mediated | (10,11,17) |

While much has been studied about the proteins and processes that mediate the transport of essential nutrients across the inner membrane very little is known about the uptake machinery that exists at the mycobacterial outer membrane. The search for OMPs is made challenging by the fact that no Msp-like proteins were identified in *M. tuberculosis*, despite the presence of over 500 homologues in other Mycobacteria and Corynebacteria (18). Bioinformatic analyses of the primary and secondary structural properties, such as the presence of repeats of amphipathic *β*-strand, N-terminal signal sequence, enrichment of cysteine residues, and the presence of a conserved C-terminal phenylalanine, were used to identify additional putative OMPs (19). A transposon library screen in *M. bovis* BCG identified CpnT (*rv3903c*), an OMP with its C-terminal domain coding for an endotoxin causing necrotic cell death. A *cpnT* mutant, similar to the *mspA* mutant, was found to have an increase in minimum inhibitory concentration (MIC), to several antibiotics (20). Enhanced glucose uptake in *cpnT* mutants has been attributed to the overexpression of SpmT (*rv0888*). Like CpnT, SpmT also possess a C-terminal sphingomyelinase domain along with a N-terminal porin forming ability, though there is currently no evidence for pore-formation. Surprising and unorthodox additions to the OMP group were α-helix-rich members of the proline-glutamate-proline (PPE) family: PPE51, characterized for glucose, glycerol and propionamide uptake, and the heme-binding surface-accessible OMPs (PPE36 and PPE62) involved in iron uptake (21,22).

Due to inherent involvement of OMPs in nutrient uptake we hypothesize that they are differentially expressed in the two distinct phases of cell growth: exponential phase is predicted to exhibit higher expression of porins involved in active uptake of nutrients, while stationary phase would allow the expression of OMPs involved in uptake of niche nutrient sources, after conventional carbon and nitrogen sources have depleted. We have extracted and enriched the outer membrane fraction of *M. tuberculosis* using a mild non-ionic detergent and employed unbiased bottom-up quantitative proteomics to comprehensively identify and compare the OMPs in both the phases of growth using a label free quantitation (LFQ) approach employing high resolution mass spectrometry. Sequence analysis allowed us to narrow our search to a few uncharacterized conserved membrane proteins and hypothetical protein. We also used NMR spectroscopy to investigate their role in uptake of small hydrophilic molecules and assessed their surface accessibility by employing immunofluorescence. We further validated their presence at the outer membrane by monitoring ethidium bromide uptake and susceptibility to hydrophilic anti-tuberculosis drugs.

## Results and discussion

### Efficient extraction of outer membrane proteins by detergents

Isolation of enriched outer membrane fraction is challenging and has impeded our understanding of the composition of the proteome of mycobacteria. We first compared the proteins extracted from exponentially growing cells using Triton X-100 and octyl-*β*-D-glucopyranoside (OBG). Since, the localization of most OMPs is not confirmed, we compared the co-extracted contaminating inner membrane and cytoplasmic proteins. In total, 1949 and 2162 proteins were extracted by OBG and Triton X-100 respectively. Large proportion of proteins (86.9%) were common between two detergents while 37 and 250 proteins were uniquely extracted by OBG and Triton X-100, respectively (Fig. S1). Known inner membrane proteins containing nucleotide binding domain or transmembrane domains (CydC, OppB, SugB, CysT, DrrB, RfbD, UgpC) were extracted by Triton X-100 but not by OBG. Due to lower abundance of established cytosolic or inner membrane proteins in OBG, it was chosen for extraction of OMPs from exponential (E) and stationary (S) phase following differential centrifugation. Samples were analyzed using tandem mass spectrometry followed by MaxQuant based analysis as described in methods. In total, 2165 protein groups were identified in 18 runs (3 technical replicates for each of the 3 biological replicates) from both the conditions. Proteins showing LFQ intensity in at least 2 of the technical replicates were averaged for each biological replicate (Fig. S2a). Around 80% of the proteins were common among the biological replicates in each condition (Fig. 2a) and LFQ intensities within each pair of biological replicates for the condition showed significant similarity (Fig. S2b). Amongst the final 1616 proteins, 74.1% (n=1198) were common while 282 and 136 proteins were present exclusively in E and S phase, respectively (Fig. 2b).

**Figure 2:**
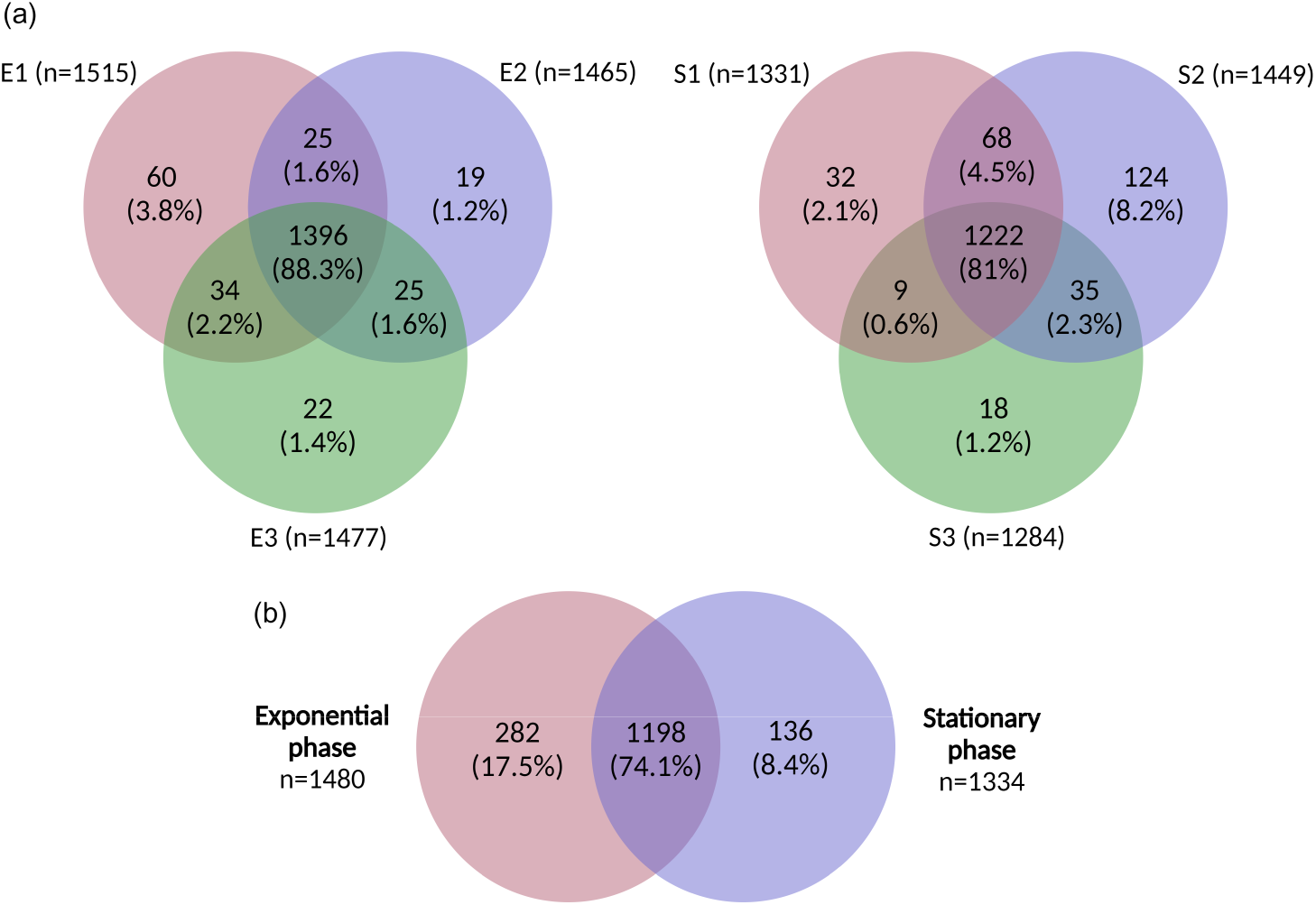
Commonality of proteins identified between (a) three biological replicates in each condition (growth phases) and (b) between the two growth phases. E, exponential phase; S, stationary phase.

Due to inherent technical challenges of specifically extracting OMPs, several criteria were used in tandem to identify contaminating proteins extracted in the OBG fraction. Though α-helix rich membrane proteins are predominant in the inner membrane, *β*-barrels have been mostly found in the outer membrane of Gram-negatives and *M. smegmatis* (23). Proteins with *α*/*β* folds are rare in membrane proteins. Prediction of transmembrane (TM) helices observed a wide distribution of TM helices amongst the extracted proteins (Fig. S3). Proteins with three and above predicted TM helices are likely to be associated with the inner membrane and thus, were discarded. Remaining 1540 proteins were found distributed across all functional classes including metabolic and information pathways (Fig. S4a). Sequence analysis along with systematic literature search were performed to ascertain the localization of proteins. A total of 614 proteins were identified as putative OMPs by analyzing data from published structural and functional studies as well as previous subcellular localization experiments. Three hundred and ninety proteins (∼24%) were previously reported in the cell wall fraction upon extraction with Triton X-114 (24) (Fig. S4b). Further, ∼20% of the proteins were identified in inner membrane or culture supernatant fractions reported earlier (25–28) (Fig. S4c). Expectedly, in this study the OBG extracted putative outer membrane proteome was dominated with proteins categorized as conserved hypotheticals and cell wall cell processes (Fig. S4a). From the functional classes observed, virulence, detoxification, adaptation, proteins showing toxin-antitoxin domain signatures were purged out while chaperones and enzymes particularly reported in terms of host pathogen interaction were retained. Channel forming proteins exist in slow growing mycobacteria since channel activity is reported in the detergent extracts of the outer membrane, however, the porins may have a different structure and oligomeric nature compared to the proteins in Gram-negatives (29).

### Proteins containing Sec/Tat signal sequence or with lipobox signatures

Since protein secretion is the prerequisite for localization of proteins to the outer membrane, presence of a signal sequence that target the protein for secretion, were considered. Most porin in Gram-negative bacteria, MspA and SpmT have a Sec type of signal that is required for translocation of unfolded proteins across the cytoplasmic membrane. We observed 72 proteins bearing Sec or Tat type of signature, 15 of them were previously found localized to the periplasm, cell surface or cell wall; although distinction between inner/outer membrane were not reported (Fig. 3a). Among the uncharacterized proteins, our study further proposes outer membrane localization of 43 proteins bearing Sec/Tat signal sequence that were not observed earlier (Fig. 3b). SignalP search annotated 72 proteins as putative lipoproteins coding for a lipobox motif, of which 10 were reported to localize to the cell surface, and 10 have a predicted substrate-binding domain. Lipoproteins LpqZ and LprC were identified in cytoplasmic fraction previously, while additional 18 proteins were reported in inner membrane fraction (30). LpqN was found to be a scaffold protein in periplasm where it binds MmpLs and interacts with cell wall enzymes (31). Interestingly, 11 of the 25 observed uncharacterized lipoproteins have not been detected upon extraction with Triton X-114 (24).

**Figure 3:**
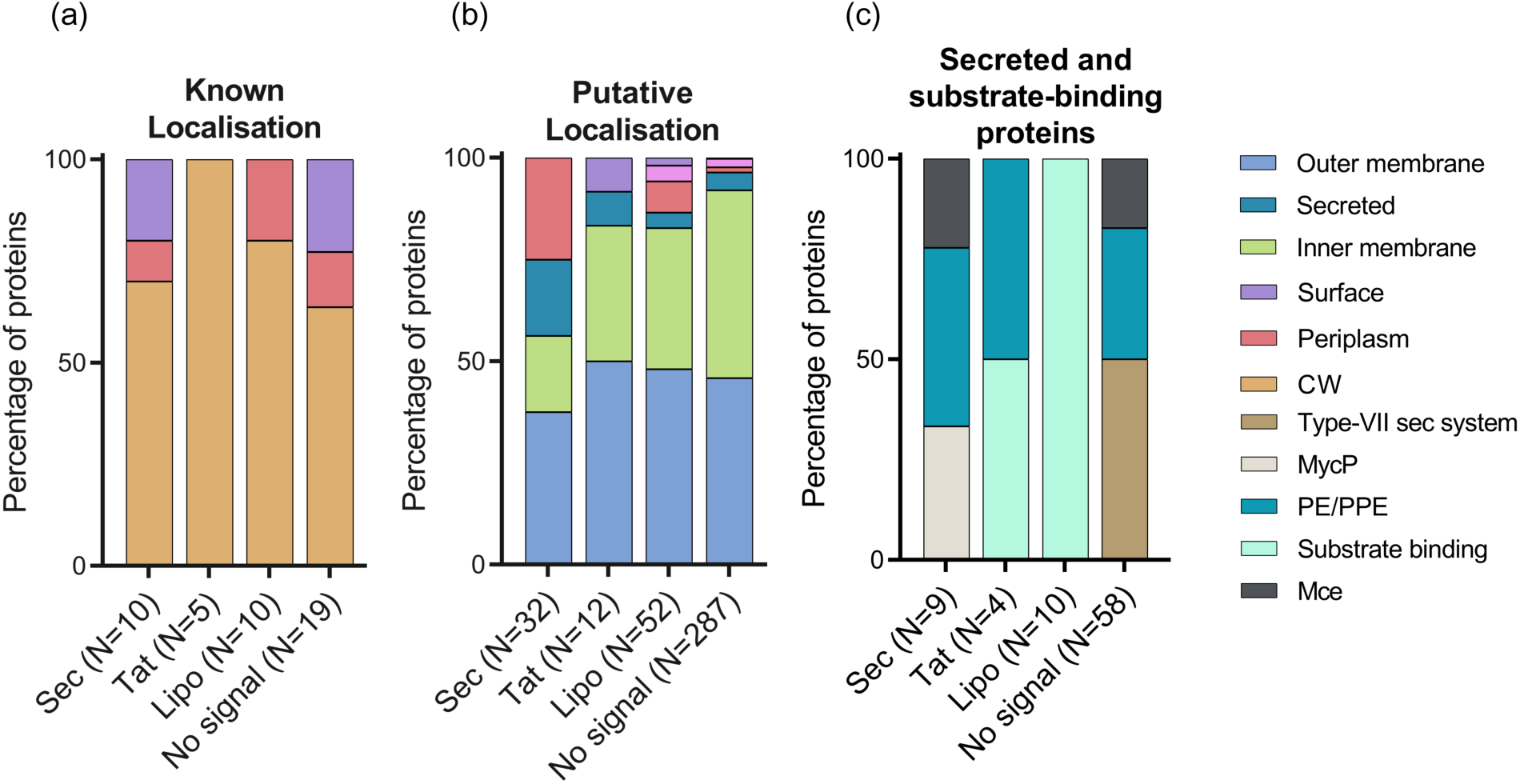
Presence of canonical signal peptides (Sec: Sec secretion signal, Tat: Tat secretion signal, Lipo: Lipoprotein signal peptide) identified using SignalP. Classification of proteins, based on localization information from previous proteomic studies and literature search, (a) Known localization (b) Putative localization (c) Secreted and substrate-binding proteins.

A large proportion of proteins lacked Sec/Tat type secretion signal sequence and 11 from this group were found to localize to the cell wall region (Fig 3a). We used clues from previous proteomic studies as well as literature to segregate remaining proteins into cytoplasmic, inner membrane or surface-localized, periplasmic or secreted (Fig. 3b, Table S1). Interestingly significant proportion of proteins were categorized as conserved hypothetical (Tuberculist; n=287) (Fig 3c). Further, 29 Type-VII secretion system proteins that lacked Sec/Tat secretion signal were identified, including secreted antigens ESAT-6 and Cfp-10 (Fig. 3c). Secreted ESX substrates (EspA/C/D/J, EsxC/D/N/O/G) were observed together with the Ecc components of the ESX-1, 2, 3 and 5. While members of the ESX-1 complex were present more in the stationary phase, other Type-VII secretion system components were overexpressed during exponential growth.

Functional class PE/PPE which occupies to 10% of the genome consists of proteins that contain PE or PPE domain in their protein sequence. Amongst 25 total proteins identified in this class, 6 proteins showed predicted secretion signal while the remaining 19 proteins lacked Sec/Tat secretion signal. *M. tuberculosis* genome bears very high number of these genes that are potentially used during pathogenesis. Functions of many of them are not yet deciphered, though recent studies have found the role of the α helical protein in nutrient uptake. Recently, PPE51 was shown to be involved in transport of hydrophilic nutrients and drugs (21). Similarly, surface accessible OMPs, PPE62 and PPE36 were proposed to be involved in heme and siderophore uptake (22) though the mechanism of heme transport across the membrane and periplasm is not understood.

In total, 12 Mce related proteins were identified and only two of them (Mce1C and Mce4A) showed predicted Sec type signal (Fig. 3c). Though no literature evidence is available for localization of Mce1C, Mce4A has been shown to localize to the cell wall fraction (32). Essentiality of Mce operons 1 and 4 for cholesterol uptake has been well established (33), although deeper structural insights in cholesterol uptake are missing and their role in the import are as yet unknown. Mycobacteria employ accessory SecA2 system for transport of proteins essential for intracellular growth, and probably uses transmembrane components of Sec translocon for export of specific proteins (34,35). Overall, we observed 222 OMPs for which no information regarding their localization could be secured, among which 162 were reported for the first time from an OBG extract of *M. tuberculosis* cell wall preparation.

### Differentially expressed proteins in Exponential and Stationary phases of growth

The final set of proteins was segregated based on their abundance in the exponential (E) and stationary (S) phase. Few proteins were identified as being unique to a particular growth phase, while 444 proteins were expressed in both E and S phases. Fold change in intensity of each protein in E phase over S phase was calculated for each pair of biological replicates and the resulting Pearson correlation coefficient values of fold change distribution ranged between 0.8 to 0.9 indicating consistency between biological replicates (Fig. S5). Based on the criterion of FC≥2, 118 and 103 proteins were found to be significantly upregulated in E and S phase, respectively (Fig. 4). Further, 123 and 47 proteins were identified as exclusively expressed in E and S phase, respectively. Peptidoglycan enzymes like PonA1, PbpA, Rv1096 and mycolic acid modulation enzymes such as Ag85 complex proteins FbpA/B/C were upregulated in E phase consistent with growth rate of growing cells. S phase was found to have a relatively higher overexpression of proteins involved in host contact and pathogenicity.

**Figure 4:**
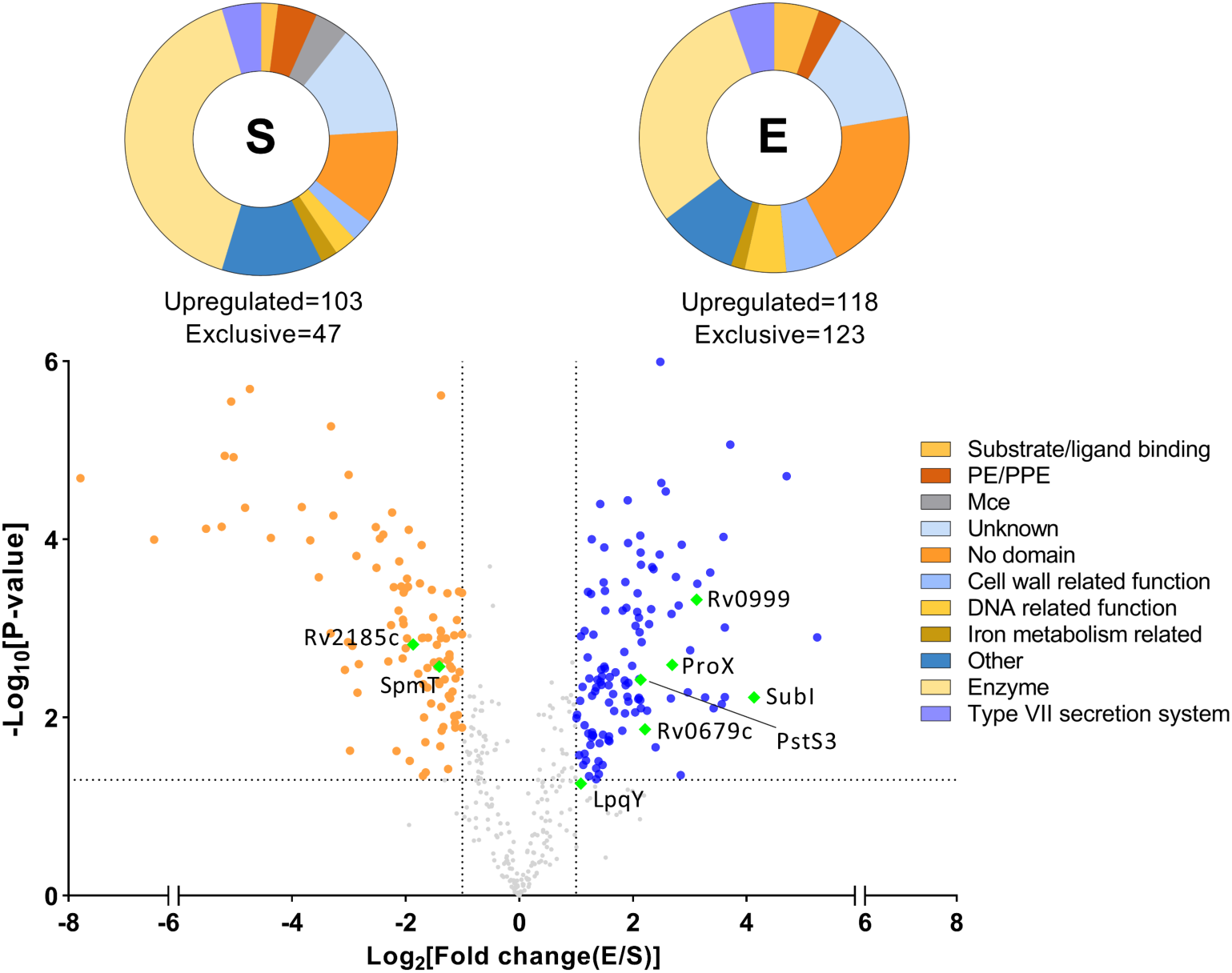
Volcano plot of proteins differentially expressed in exponential and stationary phase of growth. Significantly upregulated proteins in exponential (E) phase (blue) and stationary (S) phase (orange) with fold change >2 and p-value < 0.05. Functional classification of proteins based on protein domain. Selected representative OMPs are in green.

Interestingly, all 12 of the substrate binding proteins were found to be more in E, than in S phase (Fig S6). DppA (peptide uptake) and Rv2041c were found exclusively in E phase while SubI (sulphate uptake), ProX (betain uptake), PstS1/2/3 (phosphate uptake) and FecB (heme uptake) were significantly upregulated in E phase. Furthermore, several lipoproteins were overrepresented in E than in S phase suggesting at their role in a physiologically active cell. In contrast, S phase showed upregulation of Mce related proteins, particularly Mce1(A-D) and Mce2D. Expression of *mce1A*, *mce1D* and *mce2D* were earlier reported to be higher in stationary phase, *in vitro* in the MDR isolates (36). Out of the few PE proteins, 2 were identified exclusively in S.

Notably, PPE51, which was recently shown to be essential for uptake of small hydrophilic compounds in presence of a functional PDIM biosynthesis pathway was observed exclusively in S phase (21). Stationary phase reported higher levels of PDIMs implying that PPE51 could be an important route for entry of nutrients in stationary phase cells, though any direct evidence of its pore forming ability is still not reported (37). Importantly, higher level of PDIMs has been implicated in drug tolerance exhibited by stationary phase cells that have functional copy of PPE51 (38). This suggests that although nutrients take the route of PPE51 for cell entry, drugs may not be transported through the same route. Certain specific Esp and Esx proteins related to the type-VII secretion system showed growth specific upregulation indicating their growth rate dependent functions (Fig S6).

From the proteins represented in E and S phase, we attempted to identify proteins that can serve as transporters of antibiotics. To add an extra layer of information, conserved domains from each protein were identified using the NCBI CD database (39). Proteins were classified into distinct categories based on the presence of domains in their sequence (Fig. 4). Sixty-five proteins lacked any domain signature while an additional 54 proteins showed presence of domains with unknown function.

### Outer membrane protein variants in clinical isolates

Avoidance of drug import has been shown to be a phenotypic resistance mechanism in mycobacteria (12). Notably, the genetic locus of CpnT, a channel forming protein, was seen diversifying indicating within host positive selection by an evolutionary pressure from antibiotic exposure (20). Since whole genome sequences of clinical isolates give unique insights into the evolution of drug resistance *in vivo*, we sought to check the propensity of putative porin genes to acquire high frequency mutations in the genomes of drug resistant clinical isolates. We analysed genomic variants in publicly available whole genome sequences (WGS) of isolates that were resistant to various drugs (40–54). In order to establish a correlation of mutations with drug resistance phenotype, we quantitatively compared the mutations acquired in isolates resistant to hydrophilic drugs (test group) with that of hydrophobic drug plus pan-susceptible isolates (control group).

We identified 632 isolates from ten different genotypes with varying drug susceptibility testing (DST) profiles from 15 studies and created a dataset of isolates with known DST status. Isolates without DST data were considered susceptible for the given drug. The isolates were further segregated based on the presence/absence of known signature mutations for the individual drug (Table S2). Expectedly, known signature mutations were observed in case of well characterized drugs including, INH and RIF while most isolates lacked any marker mutation for other drugs, indicating a gap in our understanding for additional mechanism of resistance (Fig. S7a). Despite a dominance of 3 major genotypes, proportion of individual genotypes within each drug group was comparable indicating the diversity of genotypes within each drug group (Fig. S7b). A total of 1663 variants were mapped to the genomic region belonging to the outer membrane proteome, including CpnT and PPE62. Variants that were absent from both hydrophilic drug resistant isolates and singletons were discarded. The remaining 523 variants mapped to 150 putative OMP genes (Table S3). Variants that were exclusive to the hydrophilic drugs had a large proportion of SNPs and a few INDELs (Fig. S7c). Abundant amino acids in outer membrane proteome (Ala, Gly, Pro, Arg, Trp) were extensively mutated (cumulatively 41%) while low abundance amino acids (Asn, Ile, Lys, Phe, Tyr) made up 12% of the total mutated residues (Fig 5a). Most of the SNPs led to a change in chemical property of the amino acid while few instances of premature termination were observed (Fig. 5b). Also, in frame deletions were more prevalent than out of the frame deletions, while an opposite trend was observed for insertions. Large proportion of variants that were exclusive to hydrophilic drugs (n=165) were represented in 2, 3 or 4 resistant isolates cumulatively (Table S3). Due to the inequality in the number of total isolates in each drug group, we calculated frequency of resistance to each drug and calculated distribution of cumulative frequencies for each variant. Most of the probability values (75% percentile) were distributed between 0.01 to 0.04 with median value of 0.03 (Fig. S7d). We considered variants with frequency above ≥0.04 as significant variants (top 25% percentile). In order to analyse mutations that were common between 2 groups, log of odds ratio of “resistance due to mutation” over “resistance without the mutation” was calculated. The analysis revealed 48 variants that were significantly associated with resistance to hydrophilic drugs (Fisher’s exact test P<0.05, Table S3).

**Figure 5:**
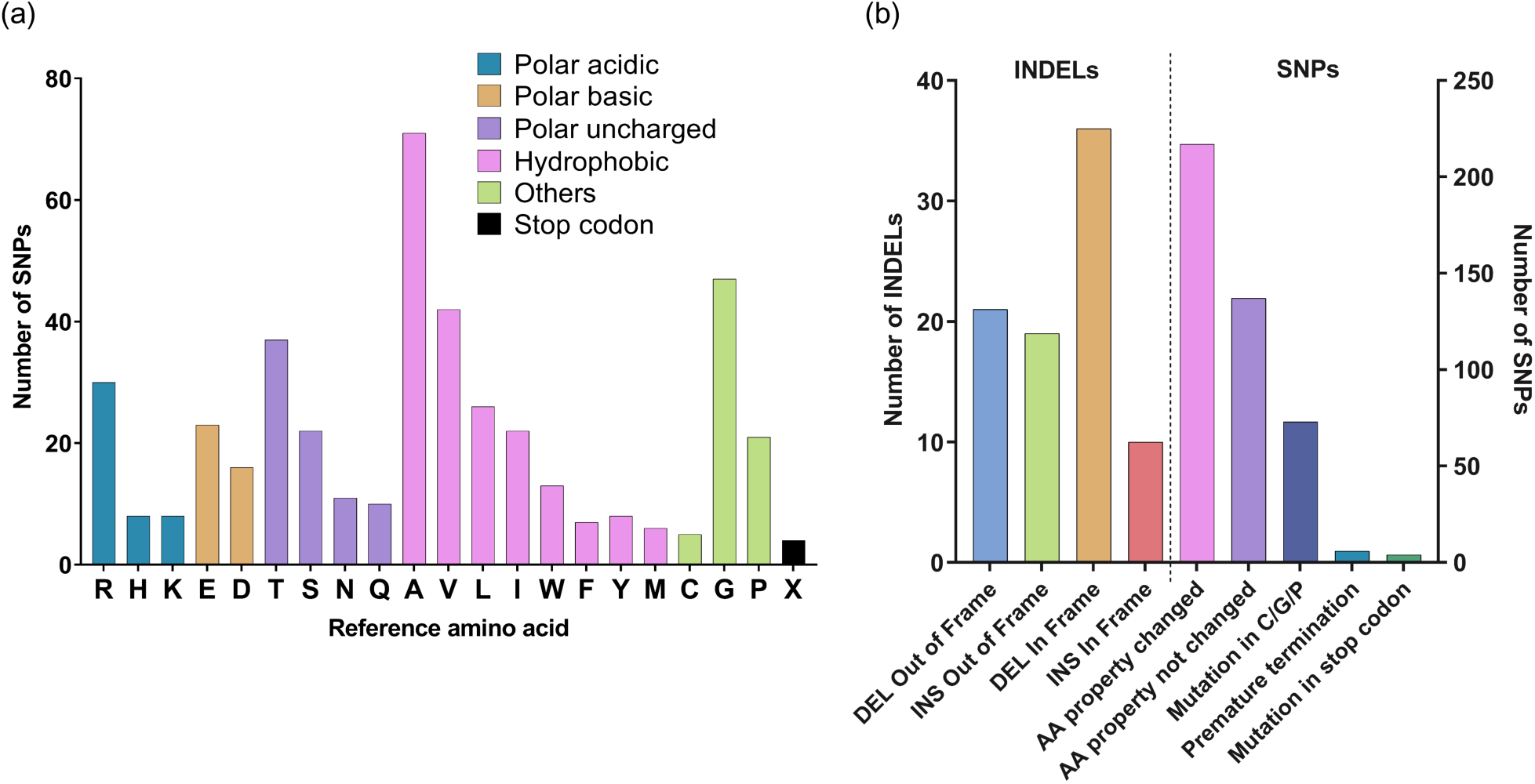
(a) Mutations in outer membrane proteome resulting in a non-synonymous SNP. Amino acids were grouped based on their chemical property. X, mutation in stop codon. (b) Number of variants plotted in each group with unique features. DEL, deletion; INS, insertion; AA, amino acid.

The variants were further mapped to genomic regions and 63 genes coding for OMPs were identified to harbour at least one of the significant variants (common and exclusive variants). Eight PEPPE genes were observed, with *ppe60* showing nine significant mutations followed by *pe-pgrs7*. *rv1192*, *pe-pgrs59* and *pe-pgrs15* recorded two INDELs each, with *rv1192* exhibiting an insertion of a base and deletion of two bases. Few additional proteins having putative functions with mycobacterial membrane were identified. *rv2717c* that forms ligand binding *β*-barrel structure showed out of frame deletion of 2 bases although mutations were restricted to isolates resistant to PZA. *rv2185c*, previously reported to be an activator of NF-*К*B, also harbours SRPBCC_2 domain structure with ligand binding pockets where a highly conserved E133 presented with an E→R substitution in the resistant isolates (55). Interestingly, *rv2585c* with an SNP (A206V) in its substrate-binding domain indicates a strong connection with resistance to hydrophilic drugs. *rv0934* (PstS1) and *rv0932* (PstS2) showed 3 and 2 SNPs, respectively, indicating an altered amino acid side chains and may influence binding to negatively charged ligands. Variations significantly associated with hydrophilic drugs gives higher fitness to the resulting mutant in presence of the drugs and represents the stepping stone mutations towards developing antibiotic resistance. This adds another layer of information about the uncharacterized proteins identified through proteomics. In the absence of structural information, the position of variation also provides clues for its importance in the molecular structures.

### Overexpression of outer membrane proteins in M. tuberculosis

Unlike Gram-negative organisms, studying mycobacterial OMPs has not been successful using sequence search for repeats of amphipathic *β*-strand leading to a *β*-barrel. Hence for *in vitro* characterization, the differentially expressed OMPs were ranked based on the following criteria: presence of a signal peptide and expression fold change higher than 4, with preference for presence of predicted substrate-binding domains and protein with conserved domain or cell wall-associated function. This resulted in prioritisation of 12 proteins for further characterization (Fig. 6).

**Figure 6:**
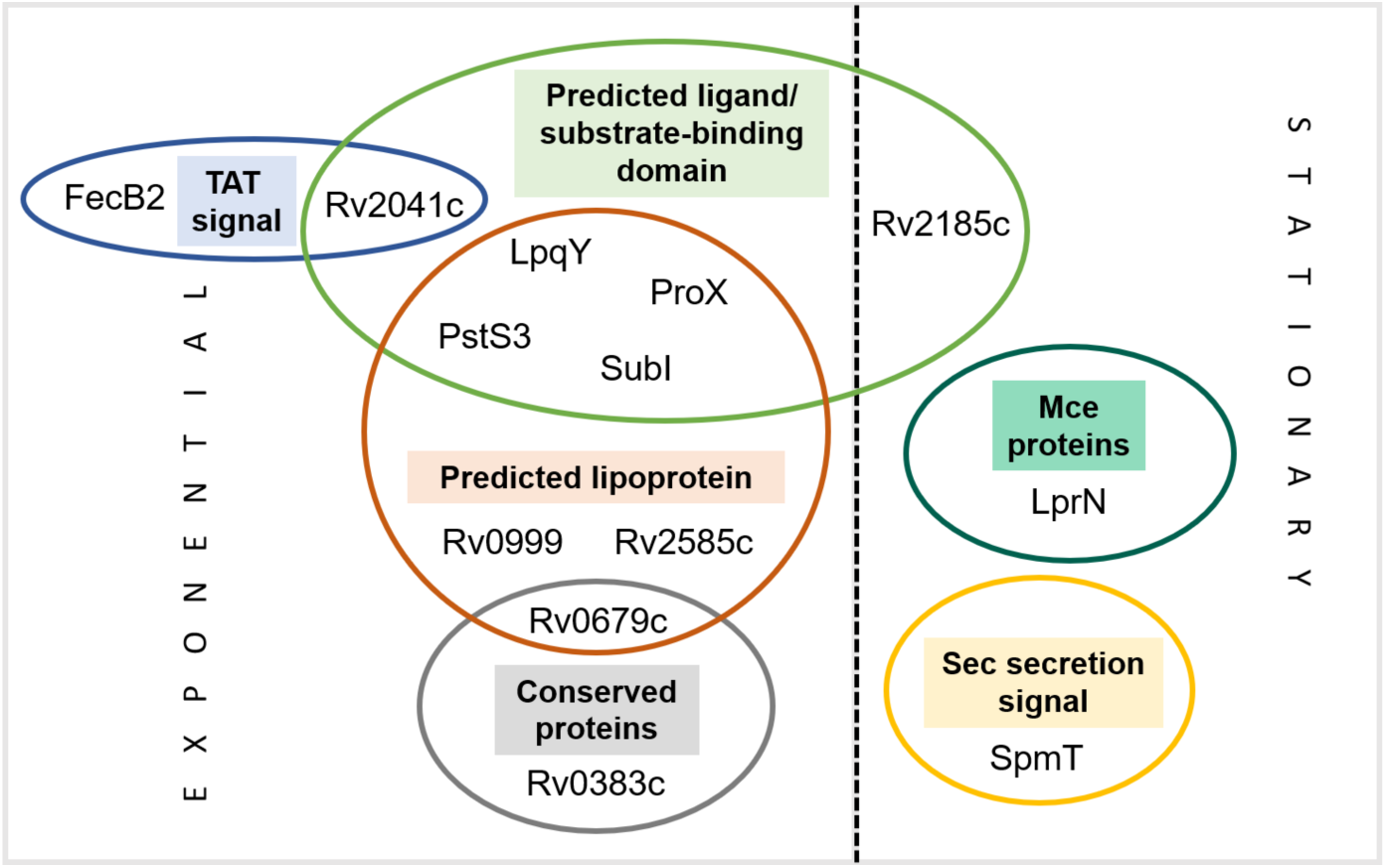
Set of differentially expressed proteins selected for characterization based on their putative outer membrane localization. Classification is based on Tuberculist.

SpmT, has been previously identified as mycobacterial OMP with a sphingomyelinase domain and served as our positive control. It was more abundant in the stationary phase along with the substrate-binding protein, Rv2185c. Rv0999, a protein with a lipoprotein signal, shows three-fold overexpression in exponential phase and its ortholog in *M. smegmatis* is structurally similar to an OMP from *Neisseria meningitidis*, FrpD (56). Proteins like FecB2, ProX, PstS1/S2/S3, LpqY and SubI are significantly overexpressed in the exponential phase and have a predicted substrate-binding domain, suggesting role in nutrient uptake (57). Rv2585c, like Rv2041c, was detected uniquely in the exponential phase with a predicted SBD. Deletion of porins has resulted in no major growth defect, suggesting the presence of other small molecule nutrient channels. Therefore, the selected OMPs were overexpressed with a C-terminal FLAG tag under a tetracycline inducible promoter (58). Protein expression was first standardized in *M. smegmatis* and induction with 200 ng/ml ATc was found to be sufficient (Fig. S8a). However, the expression was found to be leaky in *M. tuberculosis*. This may be due to the lower expression of the TetR repressor which could not suppress protein expression in the absence of inducer. However, to maintain stable protein expression, 200 ng/ml ATc was included for all experiments.

ABC system-associated lipoproteins including LpqY, SubI and ProX; TB16.3, PstS3 and FecB2 showed higher abundance in whole cell protein, while Rv2041c, SpmT and Rv2585c showed moderate expression (Fig. 7). Rv0999 could be detected only with a high-sensitive chemiluminescence substrate, suggesting relatively lower expression in the cells (Fig. S8b). Interestingly, Rv2041c and FecB2, both with Tat-secretion signal, showed an additional band for the protein with the cleaved signal sequence. The presence of plasmid overexpressing Rv0679c, a hypothetical membrane protein, was confirmed, though protein expression could not be confirmed after repeated attempts. The protein is known to be in a tight complex with the lipoarabinomannan and it is possible that the cell does not allow the protein to get overexpressed, or the protein gets cleaved at the C-terminal, removing the FLAG-tag (59). Overexpression clones of TtfA (Rv0383c) and Mce4E (Rv3495) could not be obtained even after repeated attempts. TtfA, required for the transfer of trehalose monomycolate (TMM) to the cell wall is essential in *M. tuberculosis*, though it is unclear why overexpression of the protein would not be tolerated (60). Mce4E is part of the Mce4 complex involved in transport of cholesterol across the mycolic acid layer (33).

**Figure 7:**
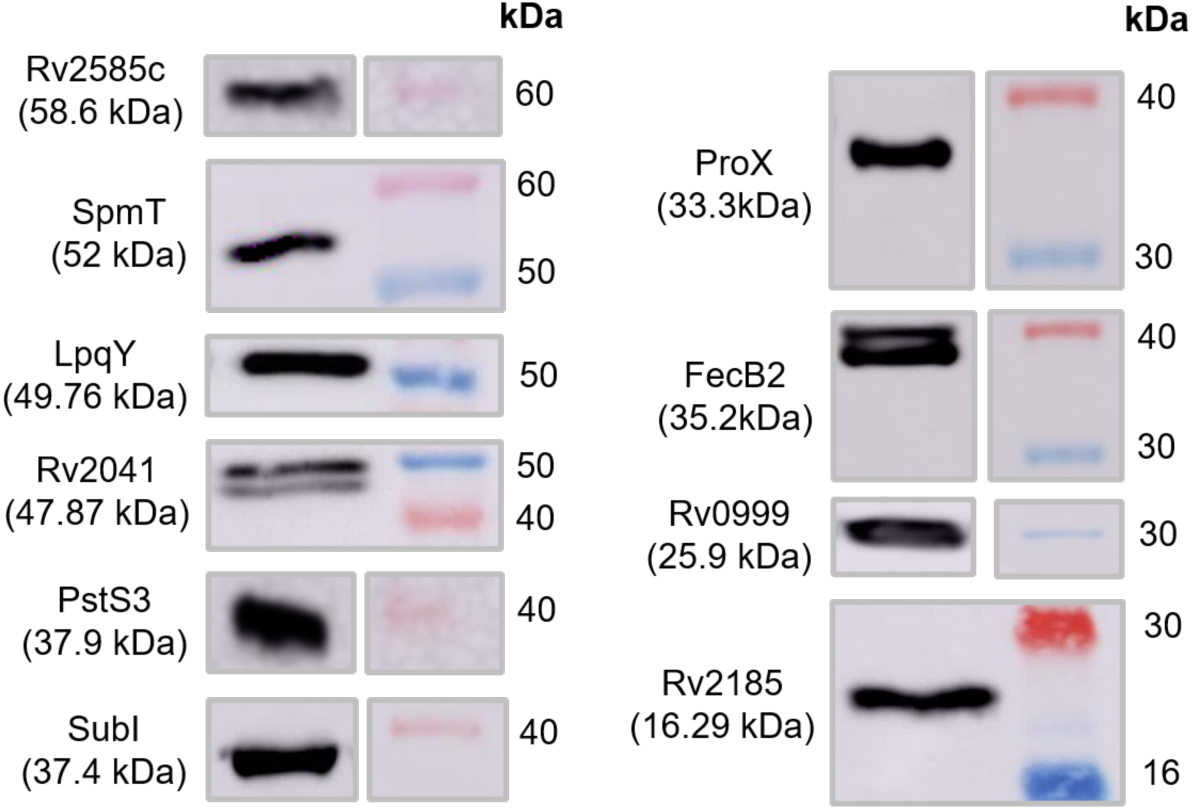
Over expression of potential outer membrane proteins in *Mycobacterium tuberculosis* mc^2^6230. Mid-exponential phase cultures were induced with 200 ng/ml anhydrotetracycline and whole cell lysates were extracted after one generation. 20 µg protein was loaded on polyacrylamide gel and probed with anti-FLAG antibody.

### Localization of the OMPs in the mycomembrane and its effect on membrane permeability

Expression of OMPs is not necessarily followed by correct localization. In order to confirm the localization to the outer membrane, differential centrifugation of the whole cell lysate and extraction with OBG was employed to enrich the outer membrane, followed by a second differential centrifugation for inner membrane and cytoplasmic fractions. Subcellular fractionation showed that Rv0999, LpqY, ProX and SubI were present in the outer membrane fraction (Fig. 8). Presence of protein in cytoplasmic fraction was expected due to constitutive protein production, while some amount of mycobacterial outer membrane that could not settle at speed of 27,000×*g* would eventually get pelleted at 100,000×*g* in the inner membrane fraction. This has been previously observed for MctB and OmpATb in *M.marinum* (61). SpmT, which was found abundantly in the stationary phase, was detected only in the whole cell lysate and further investigation showed that while it remained separated from the cytoplasmic and inner membrane fraction during differential centrifugation, it could not be solubilized by detergent. FecB2, surprisingly, was found in all fractions except outer membrane fraction. However, upon further investigation, a significant amount of protein was seen left behind in the detergent-resistant fraction of outer membrane (OBG) and inner membrane (Triton X-100) fraction.

**Figure 8:**
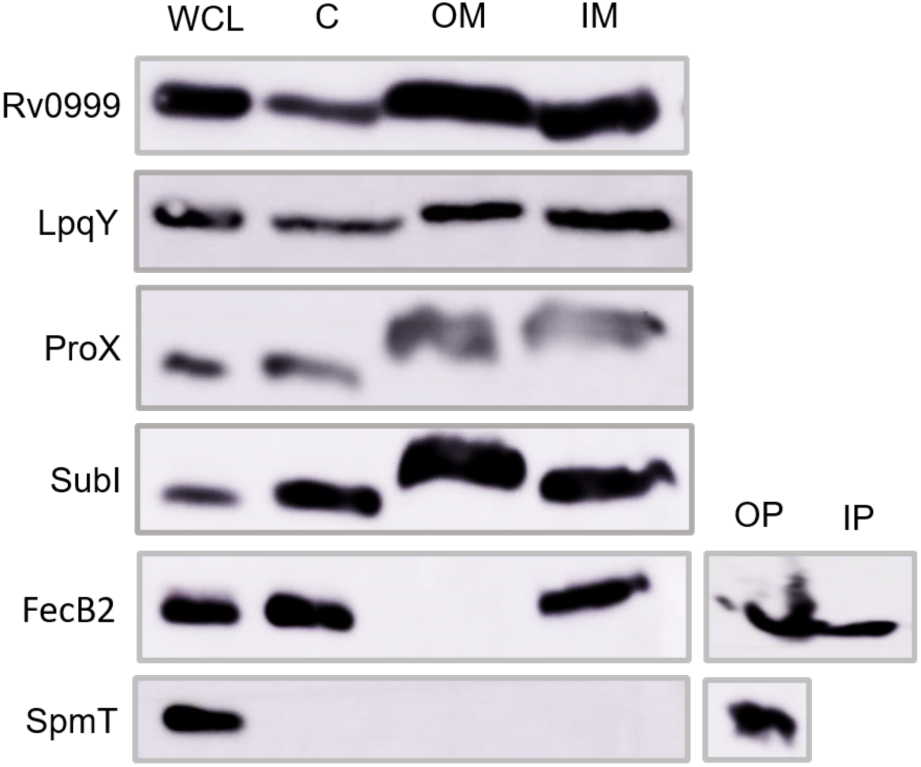
Presence of Rv0999, LpqY, ProX and SubI in outer membrane fractions of *M. tuberculosis* mc^2^6230. Rv0999 was detected using highly sensitive chemiluminescence. Equal amount (20 µg) of protein was loaded from WCL (whole cell lysate), C (cytoplasmic), OM (outer membrane), IM (inner membrane) fraction. FecB2 and SpmT were detected in detergent insoluble pellet, post extraction (OP). IP –detergent insoluble pellet of Triton X-100.

Cell surface accessibility of outer membrane proteins is what distinguishes them from inner membrane or periplasmic proteins and was assessed by probing the accessibility of the C-terminal FLAG-tag with Alexa Fluor 488-tagged antibody. Previous studies have shown that proteins present on the outer surface of cells may get stripped off due to presence of detergents (62). By growing cells in the absence of tyloxapol, we aimed to retain capsular components and the natural abundance of the surface-exposed proteins (63). While cell surface accessibility of mycobacterial OMPs has been previously shown by flow cytometry, direct microscopic evidence of their surface localization is lacking (21,64,65). A recent study reported localization of CpnT using microscopy, in its secreted state within macrophages (66). Our study for the first time specifically shows the outer membrane localization of several previously uncharacterized proteins. SpmT, a reported OMP from *M. tuberculosis*, showed significant fluorescence, corroborating with previous flow cytometry experiments, thus confirming our approach (65).

ProX and Rv2041c showed higher fluorescence among the overexpressed proteins suggesting that these proteins are surface localized (Fig. 9). ProX is a part of an ABC-binding system suggesting its outer membrane presence (67) and Rv2041c is a probable sugar-binding protein and the high fluorescence suggests association with the outer membrane as the protein also gets upregulated in presence of acid and hypoxic stress (68). Proteins like LpqY, FecB2 and PstS3 are associated with ABC complexes and show moderate fluorescence when overexpressed. LpqY is associated with the inner membrane ABC complex, SugABC and involved in uptake of free trehalose from the extracellular environment (69,70). Based on the moderate fluorescence of LpqY and its equal presence in inner and outer membrane fraction during subcellular fractionation, it is likely that the protein shuttles between inner and outer membrane. FecB2, was proposed to be involved in iron uptake and showed low fluorescence in our experiments. The protein was not detected by flow cytometry before, but is likely to be deeply embedded in outer layers of the cell envelope due its presence in detergent-resistant outer membrane fraction (22). PstS3, the most highly expressed protein in phosphate-limiting conditions was suggested to be localized to the periplasm (71). Rv2185c and Rv2585c are lipoproteins and their significant abundance in our OBG-extracted fraction and the outer membrane of *M. tuberculosis* was confirmed through immunofluorescence. SubI and Rv0999 did not exhibit fluorescence implying that while the protein is localised to the outer surface of the cell, its C-terminal end may be embedded inside.

**Figure 9:**
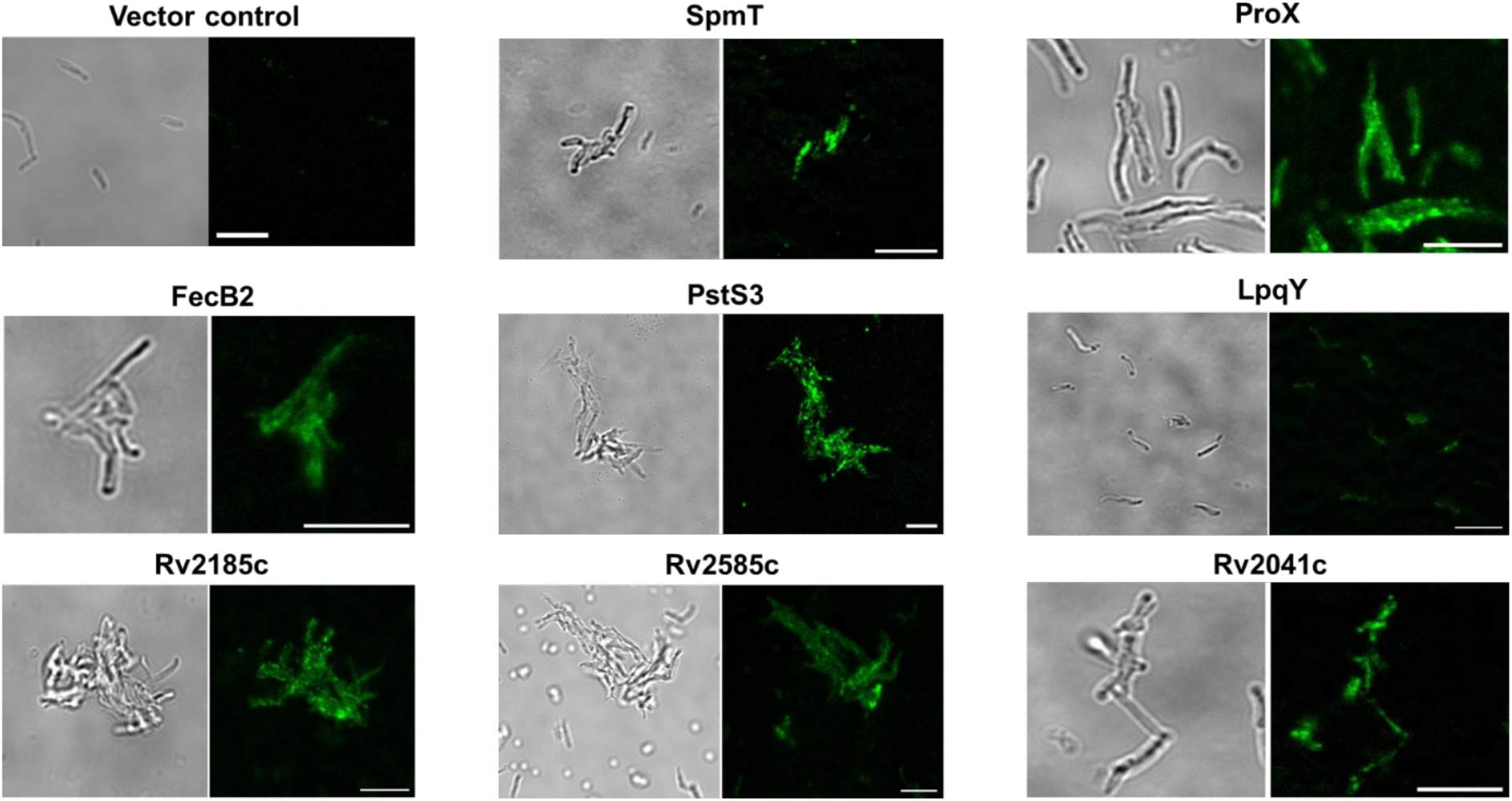
Surface localization of outer membrane proteins shown through immunofluorescence. FLAG-tagged proteins were detected using Alexa Fluor 488-labelled secondary antibody using confocal microscopy. Cells were grown without detergent and samples were processed without permeabilization. Cells overproducing SubI and Rv0999 showed negligible fluorescence. Bar = 5 µm.

Ethidium bromide (EtBr), the DNA intercalating agent, is used as an indicator of cell membrane permeability. The outer membrane serves as a barrier for EtBr permeation as spheroplasts and cells treated with outer membrane disrupting agents like polymyxin B show higher fluorescence compared to intact cells (72,73). A mycobacterium-specific study showed that the absence of major porins MspA and MspC, resulted in decreased accumulation of EtBr in *M. smegmatis* (74). Detergents used to maintain homogeneity in cultures, are found to emulsify cell wall lipids and affect its integrity. Expectedly, EtBr uptake was found to be significantly higher in the absence of tyloxapol (Fig. S9). Among the selected OMPs, overexpression of SubI, Rv0999 and FecB2 resulted in a significantly higher accumulation of EtBr fluorescence (Fig. 10). Incidentally, SubI and Rv0999 were not detected by immunofluorescence.

**Figure 10:**
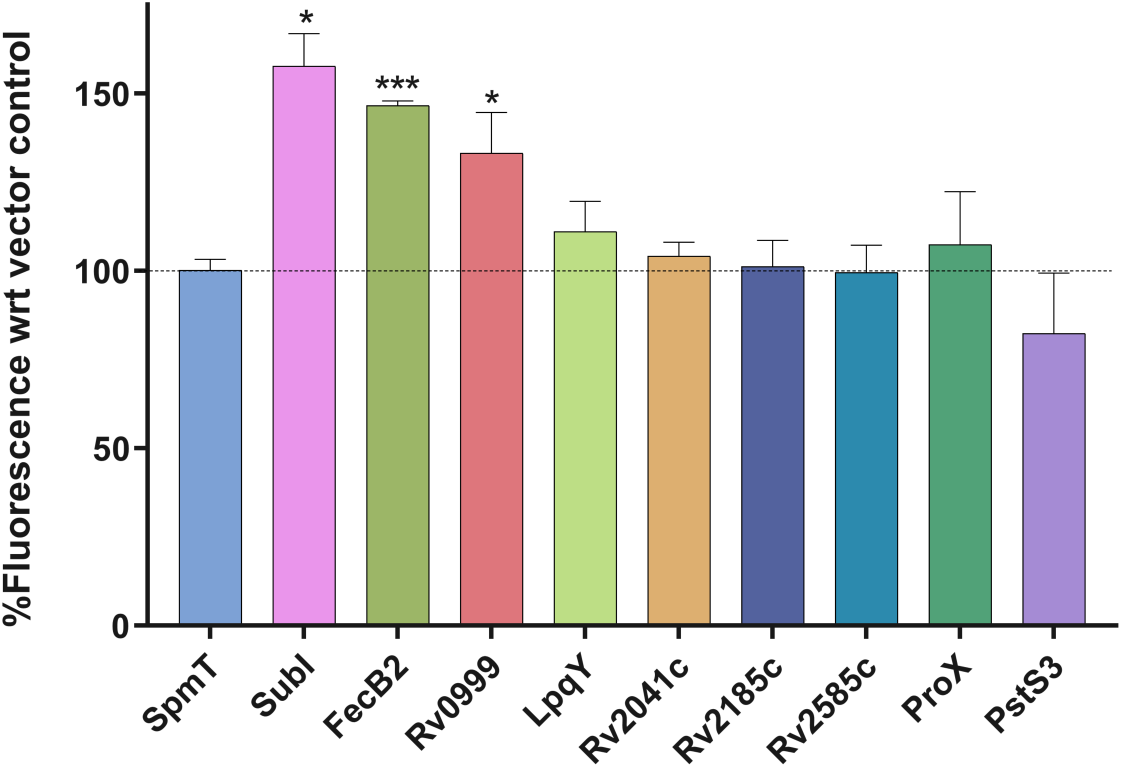
EtBr uptake assay to assess the change in membrane permeability upon overexpression of OMPs. Assay was performed in the absence of detergent and cumulative fluorescence (530/590 nm) was plotted over 15 mins, normalised against empty vector control and represented as percent fluorescence. Dotted line represents percent fluorescence for empty vector control (100%)

### Antibiotic susceptibility in M. tuberculosis overexpressing OMPs

Channel forming OMPs with a hydrophilic lumen provide ports of entry for small molecules across the thick mycomembrane. Therefore, overexpression of these proteins is expected to increase the influx for small hydrophilic molecules, making the cells hypersensitive to hydrophilic drugs resulting in a shift in the MIC. Seven antibiotics, belonging to diverse drug classes and acting through different mechanism of action, but with low log*P* values were chosen (Table 1). Small, hydrophilic drugs like isoniazid and ethambutol affects the integrity of the cell wall, while streptomycin targets protein synthesis and D-cycloserine inhibits peptidoglycan synthesis. Norfloxacin and moxifloxacin, a hydrophilic and a less hydrophilic fluoroquinolone were both tested for their uptake while rifampicin, a large hydrophobic molecule targeting the RNA polymerase was used as a control. The resazurin-based assay for measuring MIC was developed in the absence of detergents to ensure that cell surface was maintained in its native state.

While the MIC of the antibiotics remain unchanged in case of most clones overproducing the selected OMPs, clones expressing LpqY and ProX on their surface, were found to be hypersensitive for streptomycin (Fig. 11, Fig. S10). LpqY was reported to help transport the trehalose disaccharide across the periplasm to the inner membrane SugABC importer system, and thus its extracellular presence makes it an attractive candidate for drug uptake (70). Streptomycin, being an aminoglycoside is a carbohydrate derivative with a glucose and a lyxose as its constituents with multiple hydroxyl groups involved in interactions at the ligand binding site (75). Streptomycin docked to the molecular structure of LpqY with a score of –7.42. ProX, like LpqY, is also a substrate binding protein associated with an ABC transport system, ProXVWZ. Initially thought to be specific for osmoprotectants, fluorescence-quenching assay showed better interaction of ProX with polyphenols (67). Like trehalose, polyphenols with multiple hydroxyl groups, also play a role as an osmoprotectants during abiotic stress. It appears that the basis of the selectivity of ProX and LpqY for streptomycin is based on its structural similarity to trehalose. Importantly, SNPs (L184S, F247L) identified in ProX corresponds to the streptomycin binding site in the protein with a docking score of –6.11, providing further support to its role in streptomycin uptake (Fig. S11). Interestingly, PPE51-PE19 and SpmT deletion did not affect susceptibility to antibiotics and deletion of the proposed pore forming CpnT resulted in resistance to a diverse range of antibiotics including the large and hydrophobic rifampicin and clarithromycin, thus hinting at a possible indirect role in permeation through alteration of the outer membrane architecture.

**Figure 11:**
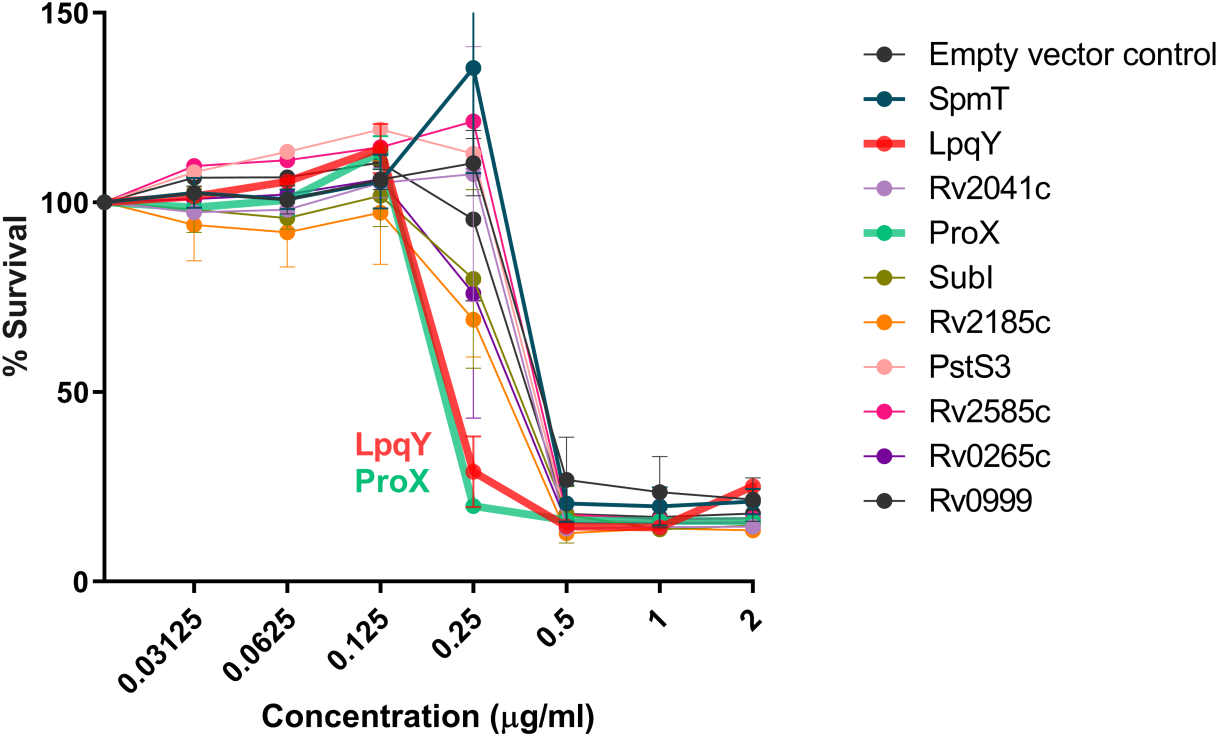
Shift in minimum inhibitory concentration (MIC) observed for streptomycin upon overexpression of LpqY and ProX. Resazurin reduction assay was performed in the absence of detergent.

### Uptake of small hydrophilic nutrients from medium

*M. tuberculosis* is a fastidious organism that requires nutrients from diverse sources and OMPs are primarily involved in nutrient uptake. While radiolabelling studies were used to probe the passage of nutrients through these proteins, they were limited by the number of small molecules that can be studied (21,64,65). We employed NMR spectroscopy as a new approach to study the nutrient preferences of *M. tuberculosis*, which can co-metabolize multiple carbon and nitrogen sources. Strains overexpressing putative OMPs were grown on a chemically defined minimal medium, with equimolar quantity of diverse carbon and nitrogen sources (0.5 mM of each glycerol, lactate, acetate, glucose, succinate, asparagine, arginine and lysine) and the spent medium was screened using 1D proton NMR in order to find the preference of the putative OMPs for a particular carbon source (Fig. 12). Multiple proteins were found to promote the uptake of glycerol. Cells overexpressing the sugar-binding proteins Rv2041c and LpqY, or the OMPs involved in the regulation of iron uptake, FecB2 and Rv0999, showed significantly high glycerol consumption. Proposed porins CpnT and PPE51 have shown their involvement in the uptake of glycerol, however, our observations suggests that the channel proteins may not be very selective for their substrate, as seen in Gram-negatives (21,64). Additionally, LpqY and FecB2 were also implicated in glucose uptake while overexpression of PstS3, a probable phosphate-binding protein, resulted in increased glucose and glycerol permeation. Overexpression of Rv2585c, a substrate-binding lipoprotein, showed increased acetate and glycerol consumption, with a preference for acetate. No discernible changes in consumption were observed in case of succinate, lactate, asparagine, lysine and arginine. NMR metabolomics allowed us to establish the connection between the OMPs and the canonical substrate that it facilitates to permeate.

**Figure 12:**
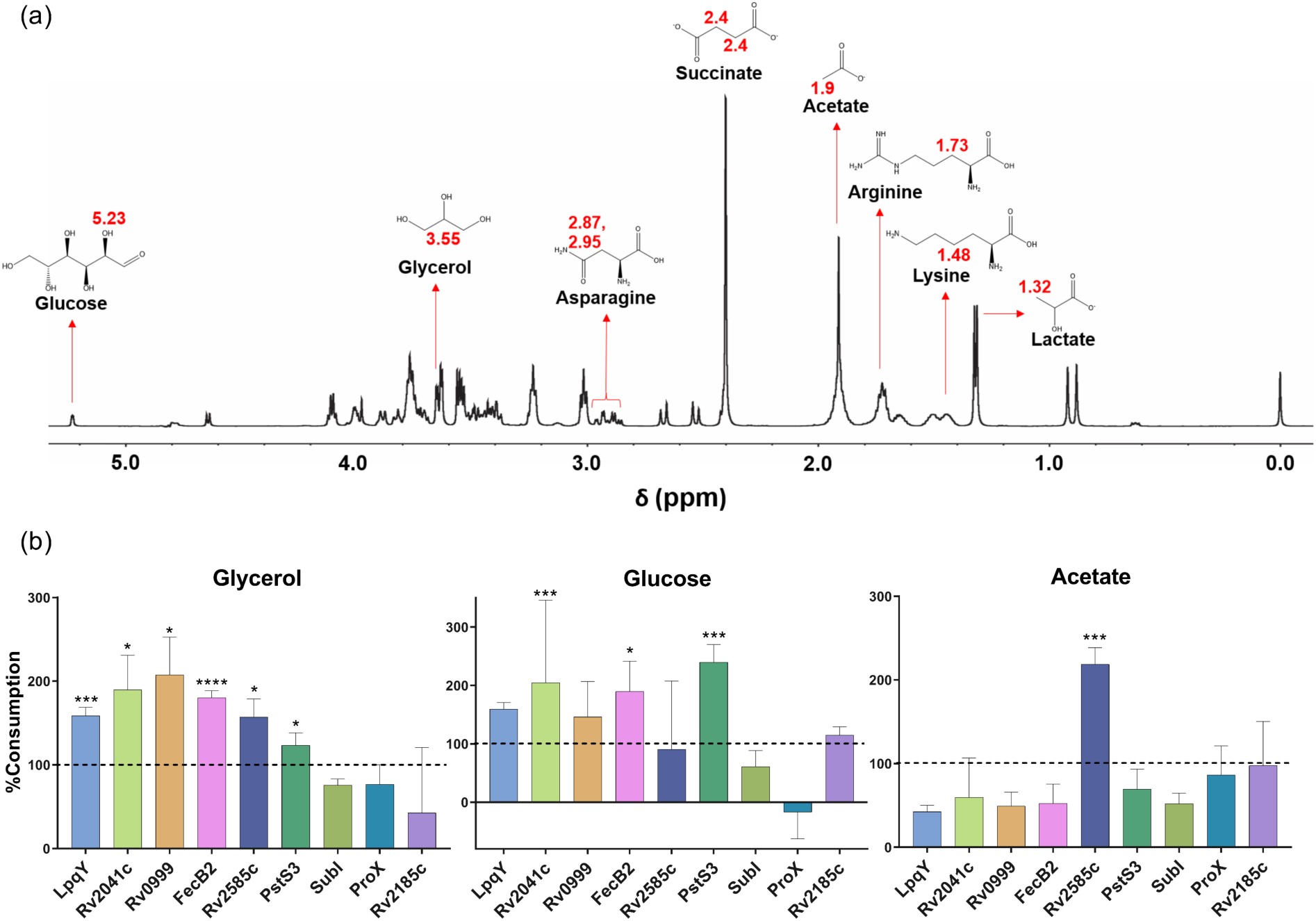
NMR metabolomics to monitor uptake of nutrients in *M. tuberculosis* strains overexpressing OMPs. (a) 1D ^1^H NMR spectrum of defined minimal medium used for monitoring nutrient uptake. Each metabolite with its unique ^1^H chemical shift, denoted in red. (b) Cells grown in defined minimal medium with 0.5 mM of each of the carbon sources. Percent consumption calculated based on the presence of each metabolite in spent medium vs control medium.

## Conclusion

The outer membrane of Mycobacteria has been an almost impenetrable barrier for molecules therefore how drugs and other molecules traverse through it is still an enigma. As we learn more about the nature and composition of mycomembrane (6,76,77), it is important to characterize the outer membrane proteome and investigate the role of the OMPs in uptake of hydrophilic molecules in *M. tuberculosis*. Identifying the pore forming proteins is a crucial step in understanding the cell envelope permeability in slow-growing mycobacteria. Knowledge of the structure and the biophysical properties of OMPs would aid in elucidating their role in drug uptake and develop effective strategies to improve permeation of drugs, which can increase the efficacy of the new and existing drugs. The aim of our study was to employ differential centrifugation, selective detergent extraction, and mass-spectrometry-based approach to comprehensively map the outer membrane proteome of *M. tuberculosis* and further validate their localization and their potential role in small molecule uptake. Knowledge of the outer membrane proteins will also help to identify the desirable surface properties in new chemical entities that should be prioritized in a critical path to increase permeability of anti-mycobacterials.

Although canonical porins have not been found in *M. tuberculosis*, uptake deficiency of small molecules have been observed in CpnT, SpmT and PPE51 depleted strains (18). Outer membrane proteins provide a passage for uptake of small hydrophilic drugs and any mutation in its internal surface can obstruct this path, resulting in antibiotic resistance. Drug susceptibility data from clinical isolates provides the best access to study genetically fixed mutations. SNPs associated with drug resistance were observed in substrate binding proteins from the genomic data reported in 15 studies and covering diverse lineages. Observed SNPs in ProX, Rv2041c and Rv2185c, in drug resistant isolates were found close to the ligand binding pocket.

ABC transporters are critical for the uptake of nutrients in mycobacteria. These inner membrane-bound complexes are associated with extra-periplasmic or surface accessible lipoproteins. LpqY and ProX, are both ABC complex-associated accessory proteins and showed hypersensitivity towards hydrophilic drugs like streptomycin, implying their role in facilitating antibiotic permeation. The proteins were abundantly present in the outer membrane fraction and their surface localisation were confirmed through immunofluorescence. LpqY has been implicated in trehalose uptake and through our NMR studies, it was also found to preferentially uptake glycerol. Together this underscores the role of the two new outer membrane protein LpqY and ProX in small molecule permeation. Mycobacterium overexpressing FecB2 and Rv0999 have also shown a higher consumption of glycerol and higher membrane permeability for EtBr. SubI, involved in sulphate uptake also showed higher EtBr accumulation.

In summary, this study documents the first ever selective extraction of OMPs in *M. tuberculosis*, and revealed potential OMPs that are surface-localized, associated with cell wall permeability and antibiotic uptake. The study also reports for the first time the surface localization of mycobacterial OMPs using immunofluorescence, and employed NMR spectroscopy to study substrate preferences. Since the surface exposed proteins were detected by immunostaining without the requirement for cell wall permeabilization, the OMPs also have good potential as vaccine candidates against *M. tuberculosis*.

## Acknowledgements

SP and AP thank IISER Tirupati for research fellowships. RM thanks SERB, Govt. of India for financial support (Grant No. ECR/2016/000665). The authors thank Prof. William R. Jacobs Jr., Albert Einstein College of Medicine, for the BSL-2 strain and Prof. Vikas Jain, IISER Bhopal, for the vector, pMTFLAG. The authors also thank Akanksha Agarwal for help with confocal microscopy and image analysis.

## Materials and Methods

### Bacterial strains, media and reagent compositions

*Mycobacterium tuberculosis H37Rv* was grown in liquid (7H9, Middlebrook) or solid (7H10, Middlebrook) media supplemented with glycerol (0.2 %), OADC (10 %). Tween-80 (0.05 %) was added only in liquid medium. *M. tuberculosis* mc^2^6230 was grown in Middlebrook 7H9 supplemented with 10% oleic acid-albumin-dextrose-catalase (OADC; HiMedia), 0.2% casamino acids (Difco), 0.5% glycerol, 24 mg/l Pantothenate (Sigma) and 0.05% tyloxapol (Sigma), in tissue culture flasks (Tarsons) in a static 37°C incubator, unless stated otherwise. *M. smegmatis* mc^2^155 was grown in 7H9 (Middlebrook) supplemented with 2% glucose and 0.05% Tween-80. Kanamycin (Sigma) was used at 20 µg/ml as a selection marker. For cloning plasmids, *Escherichia coli* was grown in Luria Bertani medium (HiMedia) with 50 µg/ml Kanamycin as the selection marker. Cultures were maintained in long term storage in glycerol stocks (30 %) at –80 °C. Compositions of PBS: NaCl (0.8%, Sigma), Na_2_HPO_4_ (0.14%, Sigma), KH_2_PO_4_ (0.024%, Sigma); Lysis buffer: PBS (50 ml), 1 tablet of complete protease inhibitory cocktail (Roche). Stock solutions of 1,4-dithiothreitol, DTT: 30 mg/ml (194 mM, Sigma) in Tris-Cl (50 mM, Sigma); iodoacetamide, IAA: 36 mg/ml (195 mM, Sigma) in Tris-Cl (50 mM). Trifluoroacetic acid (TFA), formic acid (FA) were procured from Sigma. Trypsin (sequencing grade modified) was procured from Promega. Stage tips were prepared using Empore^TM^ octadecyl C_18_ discs.

### Extraction of cell wall

Cells were grown in 3 biological replicates for exponential (E) phase upto OD_600nm_ ∼1.0 (6 generations) or stationary (S) phase up to OD_600nm_ > 5.0 (40 generations) samples. Cells were harvested by centrifugation at 5000×*g* at 4 °C and washed thrice using sterile PBS and subdivided into cell pellets such that each sample of E phase are made up of cells grown in 30 ml of culture medium while that of S phase are made up of cells grown in 4 ml of culture medium. For protein extraction, each sample was re-suspended in lysis buffer (1 ml). Cells were disrupted using bead beating of 12 cycles (40 s/cycle) at 7 m/sec (MP Biomed). Vials were kept on ice between cycles for 2 min. Glass beads were removed by centrifugation at 500×*g* at 4 °C while Unbroken cells were removed by centrifugation at 2500×*g* for 3 min at 4°C. Cell wall fraction was isolated by centrifugation at 13000×*g* for 1.5 hr at 4°C. The pellet was re-suspended in PBS (1 ml) and centrifugation was repeated at 13000×*g* for 1.5 hr at 4°C. The resulting pellet was re-suspended in octyl-*β*-D-glucopyranoside (OBG, Sigma; 1% wt/vol in PBS) or Triton X-100 (TX100, Sigma; 1 % wt/vol in PBS) and incubated for 1 hr at room temperature. The resulting insoluble matter was removed by centrifugation at 13000×*g* for 30 min at room temperature and supernatant was separated (∼1 ml). Trichloroacetic acid (1/4^th^ vol) was added to the supernatant and tube was incubated at –20 °C overnight. The pellet was recovered by centrifugation at 15000×*g* for 10 min at 4 °C. Pellet was washed with acetone (1ml) at least 3 times. The resulting pellet was solubilized in minimum volume of 6M urea for ∼30 min at 40 °C. The insoluble matter was removed by centrifugation 10000×*g* for 5 min at 4 °C and protein was estimated using BCA kit (Sigma) according to manufacturer’s instructions.

### nanoLC-MS/MS analysis

Ten µg of protein (final volume 50 µl using 6M urea) was sequentially reduced with DTT (2.5 µl) for 1 h at 37 °C, alkylated with IAA (12.5 µl) in dark for 1 h at 37 °C and again reduced with DTT (12.5 µl) for 1 h at room temperature. Subsequently, sample volume was made up to 800 µl using ammonium bicarbonate (50 mM, pH 8.0). Trypsin was added to the samples (1:5) and samples were incubated at 37 °C overnight (∼16 h). After incubation, pH was adjusted to ∼2.0 using TFA to inactivate trypsin. Samples were desalted using homemade stage tips. Briefly, peptides were bound to C_18_ stage tips previously activated and equilibrated with 100% ACN and 0.1% FA in MilliQ water (MQ) respectively. Bound peptides were washed 10 times with 0.1% FA in MQ before eluting in 50% ACN (50 µl). Samples were centrifuged at 10000×*g* for 10 min before injecting into LTQ orbitrap elite coupled with RSLC3000nanoLC (Thermo Fischer Scientific,). Samples were injected in 3 technical replicates into trap column (Acclaim PepMap^TM^ 100, 75 µm x 2 cm, nanoviper, C_18_, 3 µm, 100 Å, Thermo Fischer Scientific) and separated in analytical column (PepMap^TM^, RSLC, C_18_, 2 µm, 100 Å, 75 µm x 50 cm, Thermo Fischer Scientific) using 180 min gradient [Solvent A, 0.1% FA in water (MS grade); Solvent B, 0.1 % FA in ACN (MS grade)]. Column was equilibrated with 5% solvent B. The flow rate was 250 nl/min at 40 °C. The MS was run with automatic switching between MS and MS/MS in data dependent mode. The full MS scan was acquired in the range 350 – 2000 m/z using Fourier Transform Orbitrap (FT) operated at resolution of 60000 with activation using High Energy Collision Dissociation, automatic gain control (AGC) of 1e^6^ ions and maximum injection time of 100 ms. Each MS was followed by MS/MS of 20 most intense ions using Ion Trap (IT) with activation using Collision Induced Dissociation at 35 % normalized collision energy, AGC of 5e^3^ ions and maximum injection time of 50 msec. The minimum intensity threshold for IT was set at 500 and isolation width was 2 m/z. For MS/MS acquisition, charge state of >2 was used while charge state of 1 was rejected and a dynamic exclusion time was set at 30 sec.

### Data analysis

The raw files of all samples were analysed in Maxquant (MQ, version 1.6.2.10, (78)) using reference proteome of *M. tuberculosis H37Rv* obtained from Tuberculist ((79); version v3) and contaminants from default file present in MQ. For quantification, label free approach (LFQ) was used in MQ. Following search parameters were used: Enzyme specificity, Trypsin/P; max. missed cleavages, 2; mass tolerance for first and main search, 20 and 4.5 respectively; mass tolerance for fragment ion, 0.5 Da; fixed modification, carbamidomethylation; variable modifications, acetylation of N-terminal and oxidation of methionine; min. unique peptide required for identification, 1; min. peptide length for identification, 7; max. peptide mass, 4600 Da. Both razor and unique peptides were allowed for LFQ with min. ratio count of 2. LFQ normalization ratio was calculated using fast LFQ mode with default parameters of min. and avg. number of neighbours 3 and 6 respectively. Options of match between runs and dependent peptides were enabled. Alignment and match time windows were kept default of 20 and 0.7 min respectively. False discovery rate for all searches was kept at 1 %.

The total proteins identified in each biological replicate were defined as those having LFQ intensity in at least 2 technical replicates. Upregulated proteins in exponential and stationary phase were identified using Perseus software (version 1.6.2.2; (80)). The proteins showing valid LFQ value in only one of the technical triplicates were discarded. Further, proteins having valid LFQ in at least 2 biological replicates of both the conditions were used to draw volcano plot using fold change in LFQ intensity and p-value. The LFQ intensity value for proteins with missing value in one of the replicates was calculated by Perseus before using them for volcano plot. Proteins showing valid LFQ intensity in at least 2 biological replicates of one condition and 0/1 biological replicate in other condition were considered exclusive to the condition. Among exclusively identified proteins, proteins identified in all 3 biological replicates were regarded as high confidence while those identified in 2 biological replicates were regarded as low confidence. Rest of the proteins were discarded.

### SNP analysis

The raw files of whole genome sequences were downloaded from EMBL-ebi. The PhyResSE online workflow (version 1.0) was used for calling all variants using *Mycobacterium tuberculosis H37Rv* complete genome (NC_000962.3) as reference (81). The variants satisfying the following exclusion criteria were purged out: known variants (variants significantly associated with resistance in other studies, the list is inbuilt in PhyResSE), QUAL<500, intergenic variants, silent/synonymous mutations, total read count<20, ratio of read count of mutant allele to reference allele < 5. The remaining variants were considered for further analysis.

### Preparation of overexpression vector and cloning methods

All cloning was performed using pMTFLAG, a tetracycline-inducible shuttle vector with a pMV261 backbone and a FLAG tag at the N-terminus (58). Oligonucleotides used to amplify genes and PCR amplifying conditions are listed in Table S5. Gene sequences were amplified using NEB Phusion polymerase (New England Biolabs, USA) and ligated into EcoRV-digested pMTFLAG (New England Biolabs). Owing to blunt end ligation, plasmid was pre-treated with Alkaline phosphatase and amplicons were treated with T4 polynucleotide kinase (New England Biolabs). Cloned plasmids were subjected to colony PCR using Taq polymerase (New England Biolabs), restriction digestion analysis and Sanger sequencing in order to confirm the clones (Table S6, S7). Once confirmed, they were electroporated into *M. tuberculosis* mc^2^6230 and colony PCR was performed to confirm presence of plasmid in the mycobacteria.

### Protein expression in M. smegmatis

In order to standardize expression of proteins in mycobacterial system, the cloned plasmid for overexpression of SubI and Rv0999 was electroporated into *M. smegmatis* cells. Bacterial cultures grown to an OD_600nm_ of 0.3-0.4 were induced at 200, 300 and 500 ng/ml ATc and harvested at OD_600nm_ ∼1.2-1.5. The cell pellet was resuspended in 500 µl of 1 mM PMSF prepared in PBS and subjected to sonication: 60% amplitude; 10 cycles of 2 secs pulse ON, 30 secs pulse OFF. 10 µl of the whole cell lysate was loaded on 15% resolving gel and subjected to mouse primary anti-FLAG antibody (Thermo Scientific) and secondary goat anti-mouse HRP-conjugated antibody (Cell Signalling Technology).

### Western blot

Western blot was performed to confirm expression of proteins in *M. tuberculosis*. A secondary culture of 10 ml was grown to ∼OD_600nm_ 0.5. Protein production was induced with 50 and 400 ng/ml anhydrotetracycline (ATc), before continuing with 200 ng/ml ATc. Cells were grown under induction for one generation and homogenised using bead beater (MPBio) in lysis buffer with protease inhibitor cocktail (Roche): 12 cycles of 7 m/sec, 45 secs. Lysate was centrifuged at 10,000 rpm for 5 mins to get rid of cell debris and unbroken cells and supernatant was stored as whole cell lysate. Protein was estimated using BCA kit (Sigma Aldrich, USA) and 20 µg protein was loaded on 12% SDS-PAGE gel. Protein was transferred on nitrocellulose membrane (Biorad), probed with mouse anti-FLAG primary antibody (Thermo Scientific) and goat anti-mouse HRP-linked secondary antibody (Cell Signalling Technology, USA). Protein was visualised on Amersham Imager AI600 using Pierce ECL Western Blotting Substrate (Thermo Scientific).

### Subcellular fractionation

Subcellular fractionation was performed to ascertain presence of protein in outer membrane fraction. A large-scale (100 ml) secondary culture was grown to an OD_600nm_ of ∼0.5, when it was induced with ATc at 200 ng/ml. Cells were grown to an OD_600nm_ of 1.2-1.5 and harvested by centrifuging at 4000 rpm for 10 mins at RT. Cells were washed once with PBS and resuspended in Lysis buffer (PBS + Protease inhibitor cocktail (Roche)). The cell suspension was transferred to five bead beating tubes (Lysing Matrix B, MP Biomedicals) and subjected to lysis for 15 cycles at 7m/sec, each for 45 secs. Tubes were kept on ice for 2 mins between cycles. In order to get rid of cell debris, tubes were centrifuged at 4000 rpm for 5 mins at 4°C. A small volume of the supernatant was aliquoted and stored as whole cell lysate. Remaining supernatant was subjected to centrifugation at 27000×*g* for 30 mins at 4°C. The cell pellet was washed once with PBS and centrifugation was repeated. The pellet was then subjected to detergent extraction with 1% octyl *β*-glucopyranoside for 1 hr at RT using a tube rotisserie. The proteins extracted using detergent were obtained in the supernatant following a spin at 27000×*g* for 30 mins at RT, which is the outer membrane protein fraction. The supernatant obtained after first 27000×*g* spin was subjected to ultracentrifuge at 100000×*g* for 1 hr at 4°C. The following supernatant was stored as cytoplasmic fraction, while the pellet is the inner membrane (IM) fraction. The pellet was washed once with PBS and subjected to extraction by 1% Triton X-100 for 1 hr at RT with a tube rotisserie. The proteins extracted in the supernatant following a spin at 30000×*g* for 2 hrs at RT would be the inner membrane protein fraction.

### Immunofluorescence confocal microscopy

Cells grown to ∼OD_600nm_ 0.5 were induced at 200 ng/ml ATc and harvested after one generation. Two PBS washes ensured that cells were rid of media components. Cells were fixed with 4% paraformaldehyde (Himedia) for 10 mins, followed by 2 PBS washes. One set of cells was subjected to permeabilization with 0.1% Triton X-100 followed by 2 PBS washes. Cells resuspended in PBS (10 µl) were applied on coverslip, and air dried. 1% BSA was used as a blocking agent for 15 mins followed by 1 hour incubation with 45 µl primary mouse anti-FLAG antibody (1:1500; Thermo) and 45 min incubation with 45 µl goat anti-mouse Alexa Fluor 488-labelled secondary antibody (1:5000;). After every antibody incubation, cells were washed thrice with PBS to get rid of excess antibody. 20 µl of 5 µg/ml DAPI was added as the counterstain and incubated for 5 mins, followed by 2 PBS washes. Coverslip was inverted onto a grease-free glass slide over drops of mounting medium (80% glycerol in PBS; Glycerol, Sigma). Samples were visualised using Leica TCS SP8 Confocal Laser Scanning microscope.

### MIC assay

Minimum inhibitory concentration (MIC) assay was performed in 96-well plates with drugs belonging to different classes: Isoniazid (INH), Rifampicin (RIF), Streptomycin (STR), Ethambutol (ETM), Moxifloxacin (MOX), D-cycloserine (DCS) and Norfloxacin (NOR). A serial two-fold dilution was performed in order to obtain MIC values. Cells grown to exponential phase (OD_600nm_ ∼0.5) were diluted 1000 times to obtain OD_600nm_ 0.0005 and seeded in detergent-free medium. Cell-free control and drug-free controls were maintained. Plates were incubated at 37°C for 10 days before resazurin was added at a final concentration of 30 µg/ml. After 2 days, fluorescence readings were taken at 530/590 nm using Spectramax iD3 plate reader.

### EtBr assay

EtBr assay was used to assess cell wall permeability and was performed using a modified version of the previous protocol (74). Briefly, cells were grown to OD_600nm_ 0.8 in 7H9 complete medium without detergent. Cells once washed with PBS were pre-treated with 0.4% glucose, in order to energize the cells prior to uptake. Cells were then added to wells of black clear-bottom 96-well plate, containing EtBr solutions prepared in PBS to final concentrations 4 and 2 µg/ml. Fluorescence readings were taken at every min at 530/590 nm, for 60 mins using Spectramax iD3 instrument.

### Sample preparation for monitoring consumption of nutrients

*M.tuberculosis* cells grown in 7H9 complete medium to OD_600nm_ ∼0.8-1, were washed and subjected to nutrient starvation for 16 hrs in basal minimal medium (7.3 mM KH_2_PO_4_, 17.6 mM Na_2_HPO_4_, 190.8 µM ferric ammonium citrate, 2 mM MgSO_4_, 3.4 μM CaCl_2_, 0.35 μM ZnSO_4_, 24 mg/l Pantothenate, 0.01% tyloxapol and 20 μg/ml Kanamycin). The cells were then inoculated at a starting OD_600nm_ of 0.1 in basal minimal medium supplemented with 0.5 mM each of glucose, glycerol, acetate, succinate, lactate, asparagine, lysine and arginine. Cultures were incubated at 37°C and harvested at 36 hrs. Supernatant was concentrated 5 times and subjected to extraction with chloroform: methanol (2:1). After centrifugation at 10,000 rpm for 10 mins, upper aqueous phase was transferred to a fresh tube. A second round of extraction was performed by adding methanol: water (1:1) to the remaining organic phase and centrifuging at 10,000 rpm for 10 mins. The aqueous phases from both rounds of extraction were pooled and subjected to drying by speedvac. 600 μl of NMR buffer was added to the lyophilized *M. tuberculosis* samples. NMR buffer contains 20 mM sodium phosphate and 0.4 mM of DSS (Sodium trimethylsilylpropane sulfonate) in D_2_O (pH 7.4). 17.5 mg of DSS was dissolved in 2 ml of phosphate buffer, and then the solution was diluted (100 times) to a final concentration of 0.4 mM DSS in the buffer. Samples were homogenized with buffer using a vortex machine and then transferred to 5 mm NMR tubes for recording NMR data.

### NMR Spectroscopy and quantification of metabolites

Quad channel cryogenic probe (^1^H, ^13^C, ^15^N, ^31^P, ^2^H) installed on a AVANCE III HD Ascend Bruker NMR spectrometer with a magnetic field of 14.1 tesla was used to record all the NMR data at 298K. Noesygppr1d pulse sequence was selected from Bruker library; it is a water suppression pulse sequence and uses water presaturation in the form of RD – G_1_ – 90° –t –90° – t_m_ – G_2_ – 90° –ACQ, where RD is the relaxation delay between two successive scans of 5 sec, t is short time delay of ∼3 μsec, 90° means a 90° radio frequency pulse, t_m_ means mixing time (100 msec), and ACQ means acquisition time (6.95 s). Each spectrum was recorded for 64 scans with 16 dummy scans into a spectral width of 7200Hz and 32K data points. Parameters like water suppression, pulse width, and receiver gain were kept constant for all the ^1^H experiments recorded for different samples to avoid discrepancies. Zero filling to 64K data points and Fourier transformation were done for all ^1^H NMR spectra. Topspin (v3.5) software (www.bruker.com/bruker/topspin) was used to phase correct and baseline for all ^1^H NMR spectra manually. The singlet peak of the DSS methyl group at 0 ppm was used as a reference for the calibration of all ^1^H NMR peaks. ^1^H NMR peaks of metabolites were manually picked. Chenomx NMR suite 8.1 was used to calculate the concentration of metabolites by comparing the peaks with DSS of known concentration (0.4 mM).

