## Supplementary information for "Outer membrane proteins in *Mycobacterium tuberculosis* and its potential role in small molecule permeation"

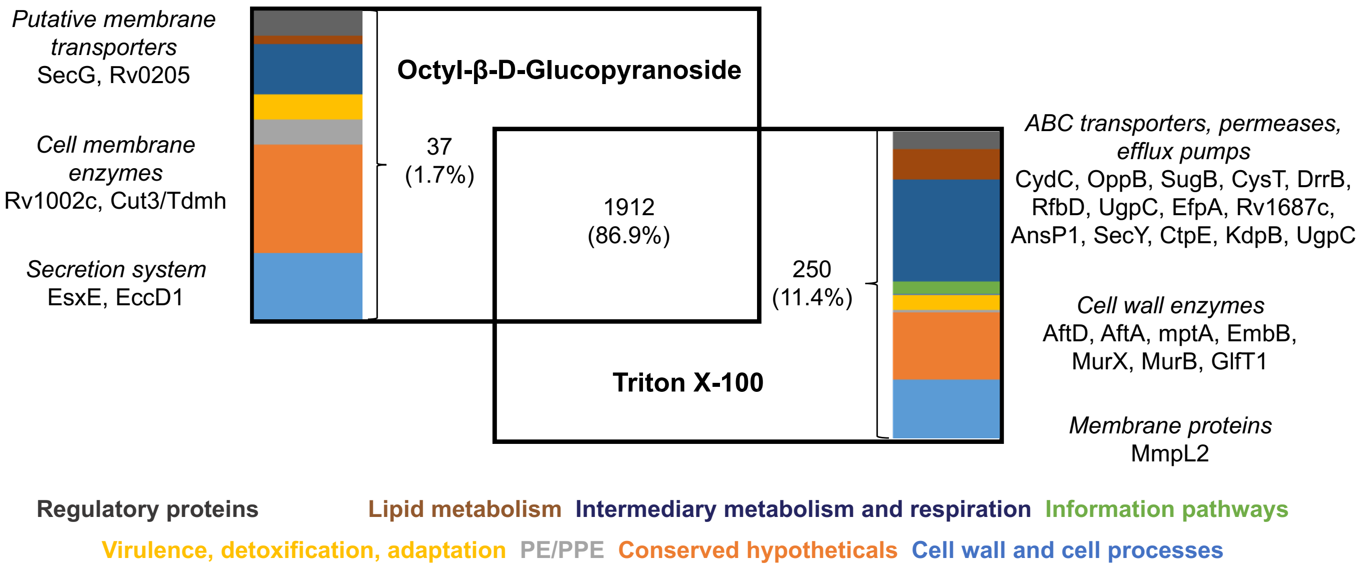


**Figure S1**: Venn diagram of proteins extracted with detergents. Proteins were extracted using octyl-$\beta$-D-glucopyranoside or Triton X-100 from 3 biological replicates of cells grown in exponential phase for each detergent. Proteins present in at least 2 biological replicates in case of each detergent were considered. Representative proteins from uniquely extracted proteins are as mentioned

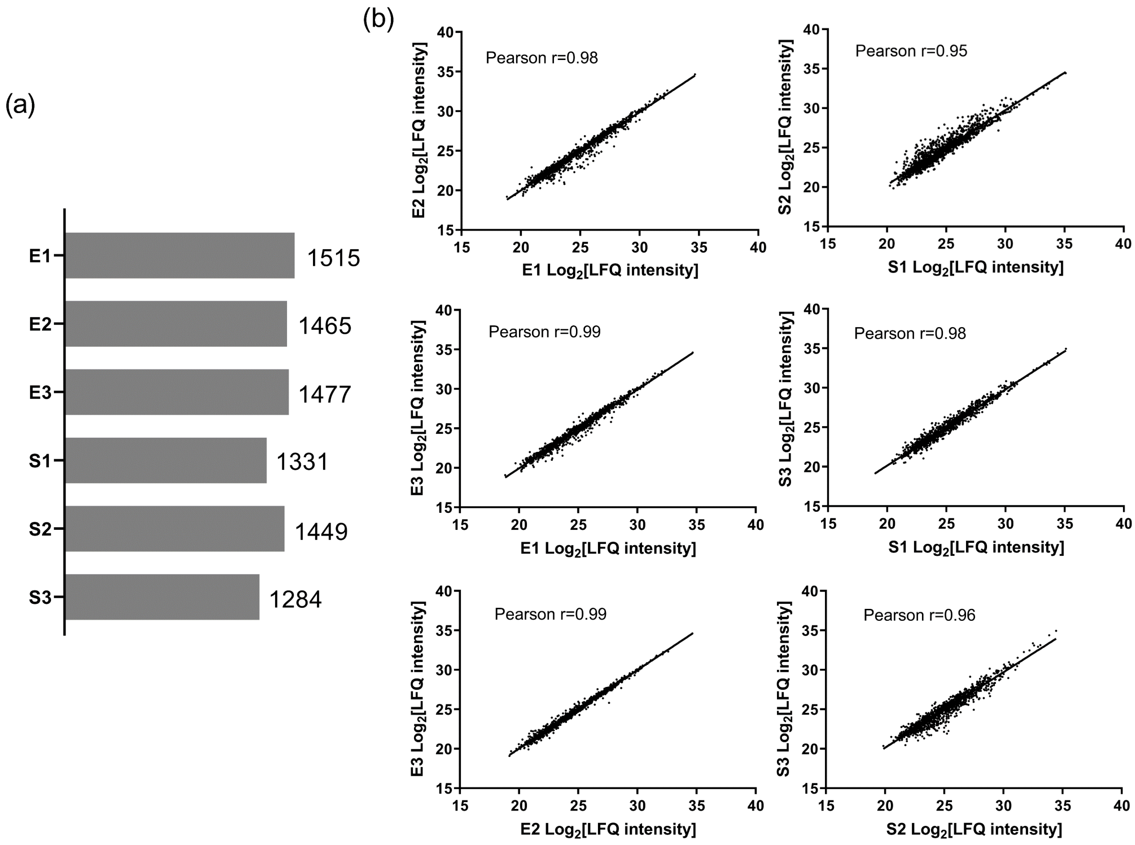


**Figure S2:** Total proteins identified in exponential phase (E) and stationary (S) phase. (a) Total number of proteins identified in each biological replicate. (b) Pairwise comparison of LFQ intensity of proteins from individual biological replicates for each condition and corresponding Pearson correlation.

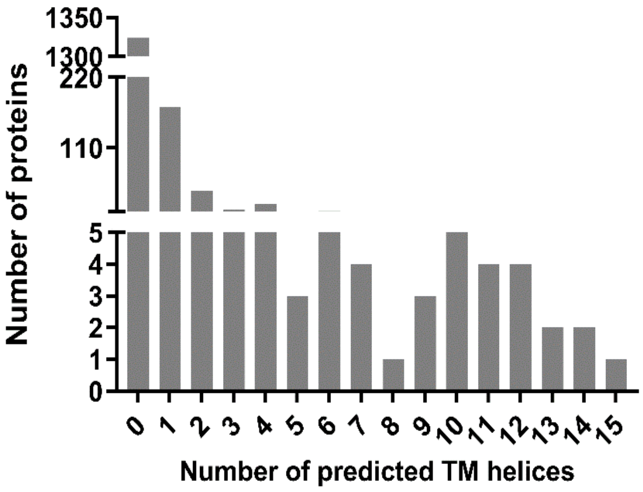


**Figure S3:** Distribution of transmembrane helices prediction using TMHMM

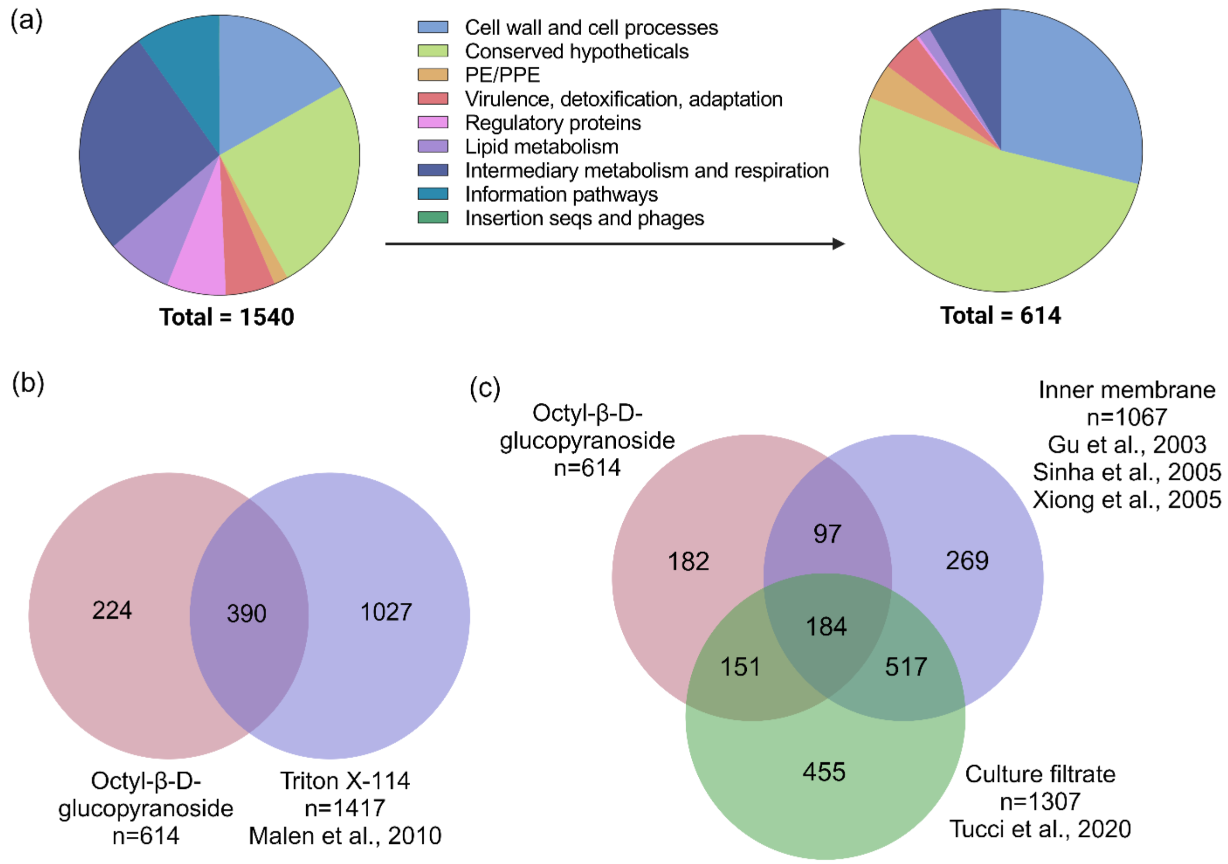


**Figure S4:** Narrowing down the search for membrane-associated proteins (a) Functional classes based on Tuberculist categories before and after removal of contaminating proteins (b) Commonality between proteins extracted in octyl-*β*-D-glucopyranoside and Triton X-114 or (c) proteins previously reported in inner membrane and culture filtrate fractions.

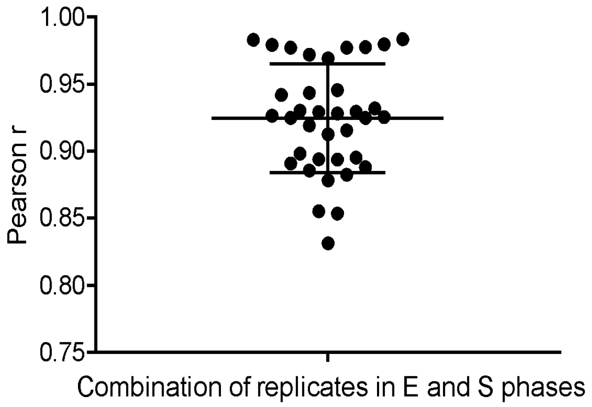


**Figure S5:** Distribution of Pearson correlation (r values) obtained from pairwise comparisons between individual biological replicates where the fold change was calculated for protein LFQ intensity in E phase over S phase.

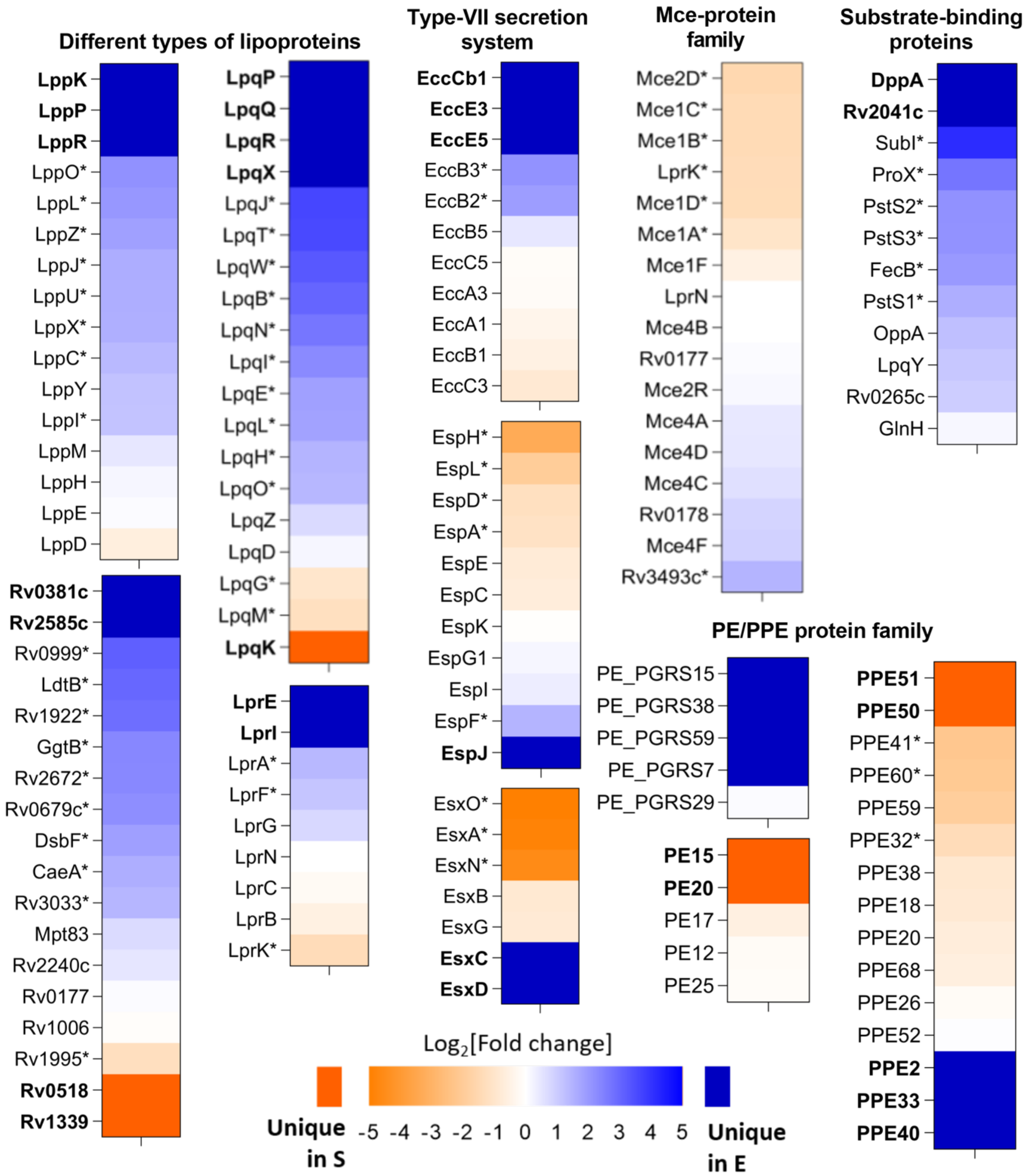


**Figure S6:** Heatmap of average fold change values showing differentially expressed substrate-binding proteins, lipoproteins, Mce proteins, PE/PPE and Esx proteins in E and S phase. Proteins in bold are uniquely present in E or S phase and * denotes statistically significant values (p-value < 0.05).

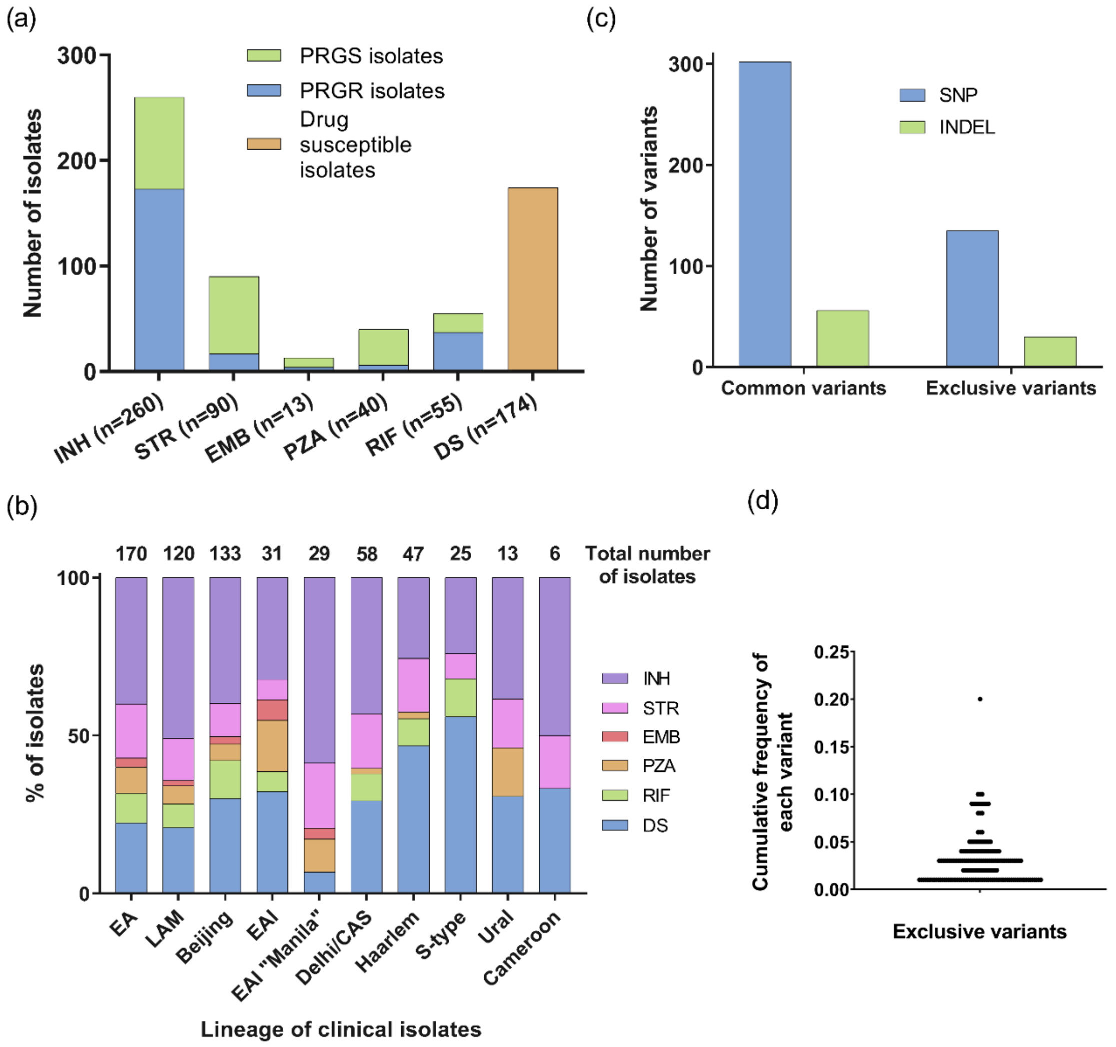


**Figure S7:** Outer membrane protein variants in clinical isolates (a) Clinical isolates showing drug susceptibility character; INH, Isoniazid; STR, streptomycin; EMB, ethambutol; PZA, pyrazinamide; RIF, rifampicin; DS, drug susceptible; PRGS- phenotypically resistant genotypically susceptible; PRGR- phenotypically resistant genotypically resistant. (b) Distribution of genotypes in the dataset of clinical isolates. (c) Total number of variants (SNP/INDEL) that were exclusive or common between 2 groups. SNP, Single nucleotide polymorphism; INDEL, insertions or deletions (d) Distribution of cumulative frequencies of individual variants that were exclusive to the hydrophilic drug resistance group.

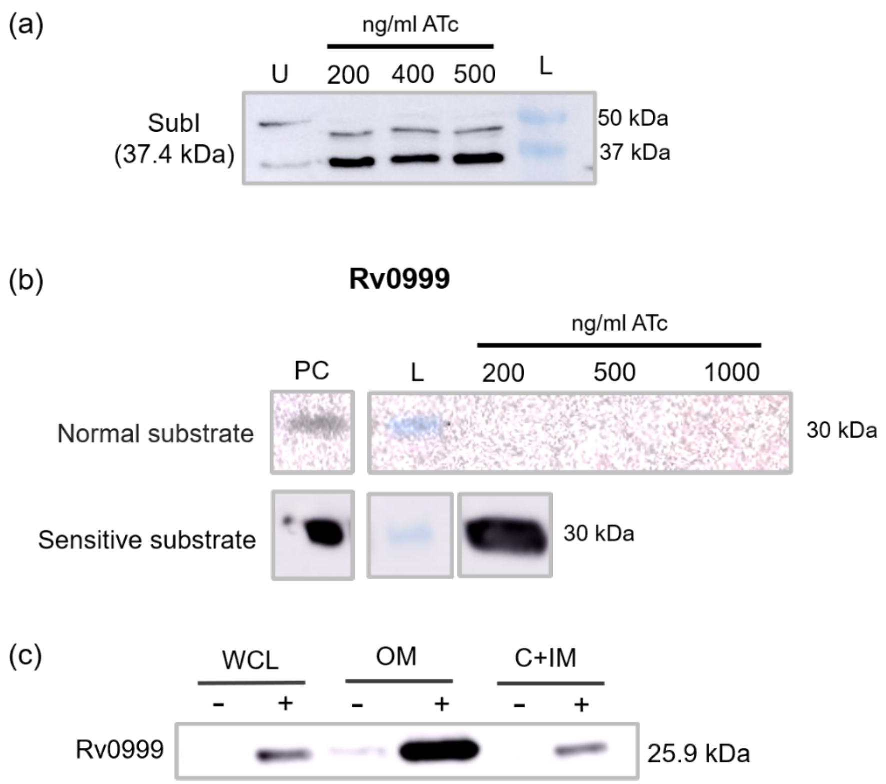


**Figure S8:** (a) Optimization of anhydrotetracycline (ATc) concentration for induction of protein expression. pMTFLAG-*subI* was introduced in *M. smegmatis* and subjected to 200, 400 and 500 ng/ml of ATc. (b) Rv0999 could be detected only using highly sensitive chemiluminescence substrate (c) Subcellular localization of Rv0999 expressed in *M. smegmatis*. 20 µg protein was loaded from WCL (whole cell lysate), OM (outer membrane), C (cytoplasmic) and IM (inner membrane), L (protein ladder).

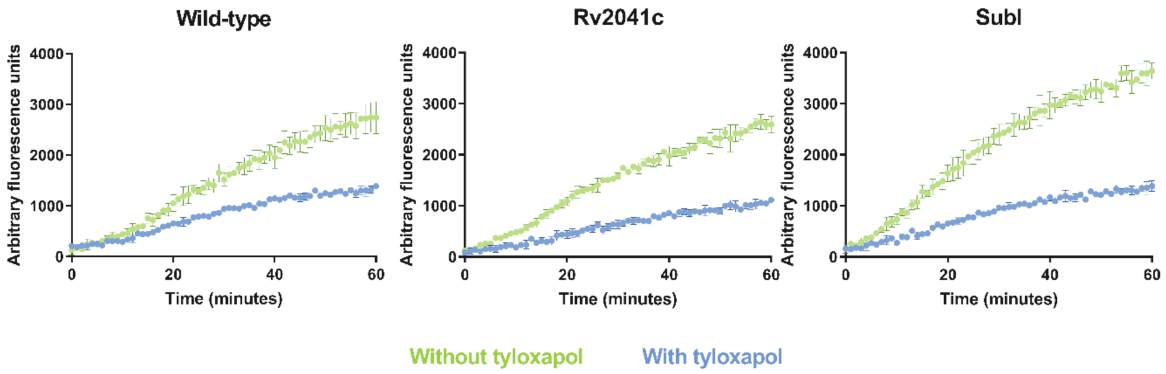


**Figure S9:** Higher fluorescence observed for cells grown and treated with EtBr without detergent. EtBr assay performed in the presence and absence of tyloxapol with 4 µg/ml EtBr. Fluorescence was monitored at 530/590 nm every minute for 60 mins.

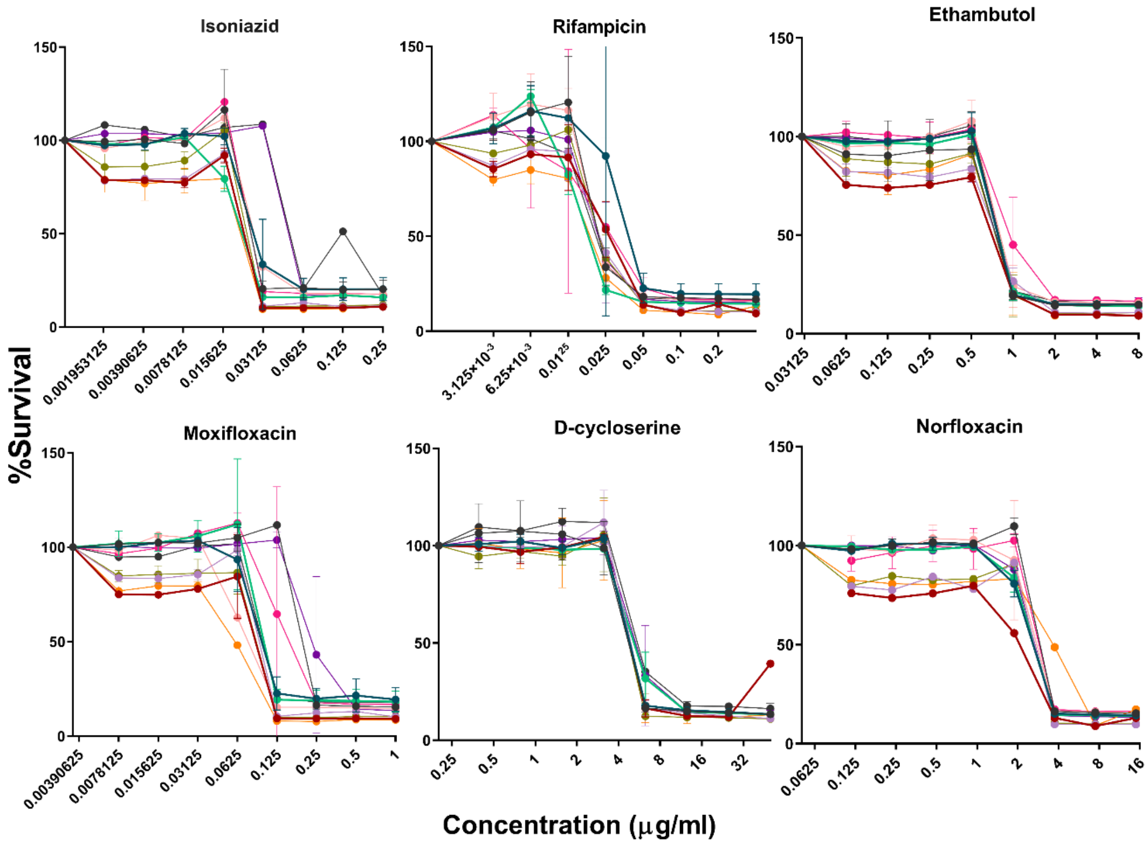


**Figure S10:** Resazurin-based MIC (minimum inhibitory concentration) assay performed, for strains overexpressing OMPs, with Isoniazid (INH), Rifampicin (RIF), Norfloxacin (NOR), Ethambutol (EMB), Moxifloxacin (MOX) and D-cycloserine (DCS). All concentrations are in µg/ml

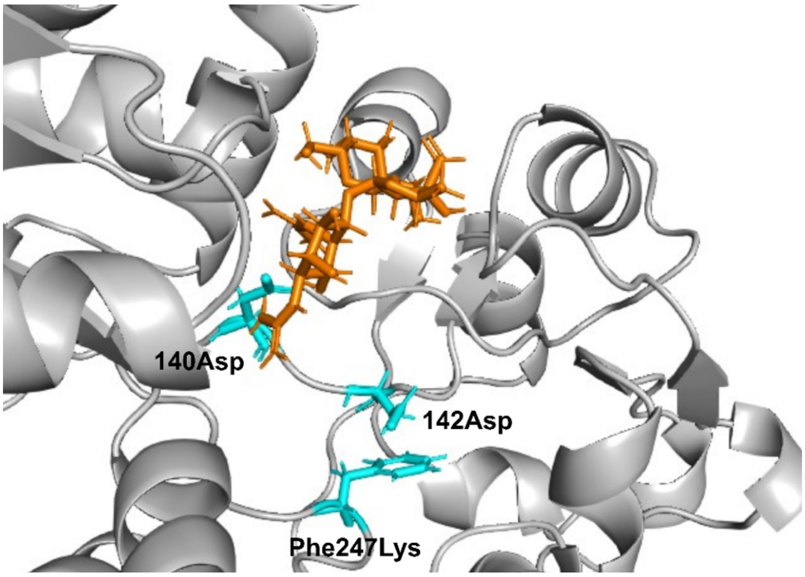


**Figure S11:** Binding of streptomycin (orange) with ProX. Substitution Phe247Lys may affect the interaction between Asp140 and streptomycin, via a loss of interaction between Asp142 and Phe247.

**Table S1: Dataset for total outer membrane proteome of *M. tuberculosis* (.xls file)**

**Table S2: Dataset from SNP analysis (.xls file)**

**Table S3: SNP variants mapping to outer membrane proteome (.xls file)**

**Table S5:** Strains and plasmids in this study

| Strains | | Source |
|---|---|---|
| H37Rv | Laboratory strain of *Mycobacterium tuberculosis* used in BSL3 facility | Dr. Amit Singh |
| $mc^2$6230 | Pantothenate auxotroph of *Mycobacterium tuberculosis* with deleted RD1 region, making it safe for BSL2 use | Prof. William R. Jacobs |
| pMTFLAG | Tetracycline-inducible replicative vector with FLAG-tag, with a pMV261 backbone | Prof. Vikas Jain |

**Table S6:** Oligonucleotides used for amplification of putative OMP genes with PCR cycle conditions using Phusion polymerase. Denat: Denaturation, Anneal: Annealing, Exten: Extension, cyc: cycles

| Gene | | Forward primer (5'-3') | Size of ampl icon (bp) | Initial denat. | Denat. | Anneal | Extn. | Final extn. | No of cyc. |
|---|---|---|---|---|---|---|---|---|---|
| Rv0265c/ FecB2 | FP | GTGCGACAGGGATGCAGCCG | 993 | 98°C 30 sec | 98°C 10 sec | 66.4°C 20 sec | 72°C 30 sec | 72°C 10 min | 35 |
| | RP | TGCGCCCAAGATCTGGCTGATC | | | | | | | |
| Rv0383c | FP | ATGGTCCCGCTTTGGTTCACGC | 855 | 98°C 30 sec | 98°C 10 sec | 63.8°C 20 sec | 72°C 25 sec | 72°C 10 min | 40 |
| | RP | GTGTTGGTAGTTGGTGGCCTGCT G | | | | | | | |
| Rv0679c | FP | GTGGTCGAGAAACCGTTGCGC | 498 | 98°C 30 sec | 98°C 10 sec | 68.3°C 20 sec | 72°C 20 sec | 72°C 10 min | 35 |
| | RP | CTCCTTGTTGATGCGGTTGCCC | | | | | | | |
| Rv0888/ SpmT | FP | ATGGATTACGCCAAACGCATCG | 1473 | 98°C 30 sec | 98°C 10 sec | 63.8°C 20 sec | 72°C 25 sec | 72°C 10 min | 40 |
| | RP | CCGTACTGCCACGTTGTCCGC | | | | | | | |
| Rv0928/ PstS3 | FP | TTGAAACTCAACCGATTTGGTGC | 1113 | 95°C 3 min | 95°C 30 sec | 65.3°C 20 sec | 72°C 45 sec | 72°C 10 min | 35 |
| | RP | GGCGATTGCTTTGACCGCAGTC | | | | | | | |
| Rv0999 | FP | GTGCGTCCACCACTGGCACCG | 759 | 98°C 30 sec | 98°C 10 sec | 69.5°C 20 sec | 72°C 20 sec | 72°C 10 min | 40 |
| | RP | ACCCCGTACTGCTGCTACCGCCT GTAC | | | | | | | |
| Rv1235/ LpqY | FP | GTGGTCATGAGTCGCGGGCG | 1407 | 98°C 30 sec | 98°C 10 sec | 66.4°C 20 sec | 72°C 35 sec | 72°C 10 min | 35 |
| | RP | CGGGAGCAGACCCATGCCGTC | | | | | | | |
| Rv2041c | FP | ATGGTCAATAAGCCGTTCGAGCG G | 1320 | 98°C 30 sec | 98°C 10 sec | 66.4°C 20 sec | 72°C 35 | 72°C 10 min | 40 |

| | | | | | | | | | |
|---|---|---|---|---|---|---|---|---|---|
| | RP | TGGATTTCGCAGCACTTCATCGAC | | | | | sec | | |
| Rv2185c/ TB16.3 | FP | GTGGCGGACAAGACGACACAG | 435 | 98°C 30 sec | 98°C 10 sec | 66.4°C 20 sec | 72°C 20 sec | 72°C 10 min | 35 |
| | RP | GCCCTCGACTCGTTTCTTCAGATC | | | | | | | |
| Rv2400c/ SubI | FP | ATGCTCTCCTTGACGCTTTCTG | 1071 | 98°C 1 min | 98°C 10 sec | 65.3°C 20 sec | 72°C 20 sec | 72°C 10 min | 35 |
| | RP | TCCGGTGGCCCGCAGATAAATC | | | | | | | |
| Rv2585c | FP | GTGGCACCACGACGACGGCGTCA TA | 1674 | 98°C 30 sec | 98°C 10 sec | 66.4°C 20 sec | 72°C 35 sec | 72°C 10 min | 35 |
| | RP | CCGAGCCAGAGCCCAGCGATC | | | | | | | |
| Rv3495 | FP | ATGAACCGAATCTGGTTGCGC | 1155 | 98°C 30 sec | 98°C 10 sec | 66.4°C 20 sec | 72°C 30 sec | 72°C 10 min | 35 |
| | RP | **CTGTCCCGATGCCGTACCGG** | | | | | | | |
| Rv3759c/ ProX | FP | ATGAGGATGCTGCGACGCCTAC | 948 | 98°C 30 sec | 98°C 10 sec | 65.3°C 20 sec | 72°C 20 sec | 72°C 10 min | 35 |
| | RP | CTGCCGCACTGGATGATCGAAAC C | | | | | | | |

**Table S7:** Oligonucleotides used for confirmation of presence and orientation of amplicon after blunt end cloning in pMTFLAG

| **Primer name** | **Forward primer (5'-3')** | |
|---|---|---|
| pMTFLAGF | GACGTGCGGTCGAGACGCTTC | |
| **Gene** | **Reverse primer (5'-3')** | **Size of amplicon** |
| Rv0265c | CTG ATA GGT ATC GGC GTC CAC C | 572 |
| Rv0383c | CCA GAT GTC GCT TTC TCG TGG C | 521 |
| Rv0679c | CGT CGA TCC TGT CGA AGG CGA TC | 552 |
| Rv0888 | GGT TGC CAT CGG GGT CGC AG | 561 |
| Rv0928 | CAC CGG CAG ATT CCA CGC CG | 539 |
| Rv0999 | CCA TGT TGC CCT GCA ACC CG | 534 |
| Rv1235 | GTA CAG CTT GTG GTT CCA GCC G | 587 |
| Rv2041c | CGG AAA GGC GTA CTG GCC TC | 602 |
| Rv2185c | GCT TGA GCA TCC CGA TCA TGG | 525 |
| Rv2400c | CCC TTG GTG GCA TCG GCG TC | 571 |
| Rv2585c | CTG CGT AGC AGC GTC GAA GC | 578 |
| Rv3495 | CGA GCC GTC AAC CAA CCT CC | 557 |
| Rv3759c | GTC AGG ATC GAC AGA TCG CCG | 556 |

**Table S8:** Details of restriction enzymes used for confirmation of cloned plasmids and expected fragments

| Gene | Size of cloned plasmid (bp) | Linearization | | Double digestion | | Expected fragments | Expected fragments with incorrect orientation | Expected fragment with plasmid control |
|---|---|---|---|---|---|---|---|---|
| | | Enzyme | Buffer | Enzyme | Buffer | | | |
| Rv0265c/ FecB2 | 5799 | BamHI | 3.1 | BglII, BamHI | 3.1 | 2955 bp, 1844 bp, 1000 bp | 3923 bp, 1844 bp, 32 bp | 2965 bp, 1844 bp |
| Rv0383c/ TtfA | 5661 | BamHI | 3.1 | BamHI | 3.1 | 4924 bp, 737 bp | 5504 bp, 157 bp | 4809 bp |
| Rv0679c | 5307 | BglII | 3.1 | PstI, EcoRI | CutSmart | 4564 bp, 740 bp | 4265 bp, 1039 bp | 4809 bp |
| Rv0888/ SpmT | 6279 | KpnI | 1.1 | SacI | 3.1 | 3810 bp, 1019 bp, 925 bp, 525 bp | 2560 bp, 2175 bp, 1019 bp, 525 bp | 3265 bp, 1019 bp, 525 bp |
| Rv0928/ PstS3 | 5919 | BamHI | 3.1 | SmaI | CutSmart | 2822 bp, 2118 bp, 979 bp | 2500 bp, 2118 bp, 1301 bp | 2691 bp, 2118 bp |
| Rv0999 | 5565 | BamHI | 3.1 | BglII, SalI-HF | 3.1 | 3644 bp, 1921 bp | 3000 bp, 2565 bp | 4809 bp |
| Rv1235/ LpqY | 6216 | BamHI | 3.1 | KpnI | 1.1 | 5015 bp, 1198 bp | 4549 bp, 1664 bp | 4809 bp |
| Rv2041c | 6126 | BglII | 3.1 | BamHI | 3.1 | 1275 bp, 4851 bp | 84 bp, 6042 bp | 4809 bp |
| Rv2185c | 5241 | BglII | 3.1 | BglII, SalI-HF | 3.1 | 3040 bp, 2201 bp | 3275 bp, 1966 bp | 4809 bp |
| Rv2400c/ SubI | 5877 | BglII | 3.1 | BamHI, PvuII | 3.1 | 4842 bp, 1028 bp, 7 bp | 5795 bp, 75 bp, 7 bp | 4809 bp |
| Rv2585c | 6483 | NdeI | 3.1 | EcoRV-HF, KpnI-HF | CutSmart | 4923 bp, 783 bp, 774 bp | 4134 bp, 1572 bp, 774 bp | 4080 bp, 729 bp |
| Rv3495/ LprN | 5961 | NotI-HF | CutSmart | PstI-HF, NotI-HF | CutSmart | 4333 bp, 1628 bp | 4963 bp, 998 bp | 4809 bp |
| Rv3759c/ ProX | 5754 | BamHI | 3.1 | EcoRI | Buffer EcoRI | 4342 bp, 1412 bp | 4937 bp, 817 bp | 4809 bp |
